# Dual process impairments in reinforcement learning and working memory systems underlie learning deficits in physiological anxiety

**DOI:** 10.1101/2025.02.14.638024

**Authors:** Jennifer Senta, Sonia Bishop, Anne GE Collins

## Abstract

Anxiety has been robustly linked to deficits in frontal executive function including working memory (WM) and attentional control processes. However, although anxiety has also been associated with impaired performance on learning tasks, computational investigations of reinforcement learning (RL) impairment in anxiety have yielded mixed results. WM processes are known to contribute to learning behavior in parallel to RL processes and to modulate the effective learning rate as a function of load. However, WM processes have typically not been modeled in investigations of anxiety and RL. In the current study, we leveraged an experimental paradigm (RLWM) which manipulates the relative contributions of WM and RL processes in a reinforcement learning and retention task using multiple stimulus set sizes. Using a computational model of interactive RL and WM processes, we investigated whether individual differences in physiological or cognitive anxiety impacted task performance via deficits in RL or WM. Elevated physiological, but not cognitive, anxiety scores were strongly associated with worse performance during learning and retention testing across all set sizes. Computationally, higher physiological anxiety scores were significantly related to reduced learning rate and increased rate of WM decay. To highlight the importance of modeling WM contributions to learning, we considered the effect of fitting RL models without WM modules to the data. Here we found that reduced learning performance for higher physiological anxiety was at least partially misattributed to stochastic decision noise in 9 out of 10 RL-only models considered. These findings reveal a dual-process impairment in learning in anxiety that is linked to a more physiological than cognitive anxiety phenotype. More broadly, this work also points to the importance of accounting for the contribution of WM to RL when investigating psychopathology-related deficits in learning.

## Introduction

Our ability to learn from our experiences of the world is a crucial element of successful decision-making and ultimately survival. Along with other dimensions of psychopathology, anxiety has been behaviorally linked to impairments in learning, including slower learning and reduced performance (1). Models of reinforcement learning (RL; (2) have been successfully used to investigate the cognitive mechanisms of learning across animals and humans. Extending this work into the clinical domain, RL models have been used to investigate the influence of psychopathology upon learning (3). Here, there remains a lack of clarity regarding the precise effects of anxiety on the computational mechanisms supporting reinforcement learning (4). This is complicated by differences between studies in both the experimental paradigms used (5–7) and in whether anxiety symptoms are directly measured or if a stressor manipulation is used as a proxy for anxiety (4,8). In relation to the latter, prior work suggests that individual differences in anxiety-related psychopathology and induced anxiety may have very different neural signatures (9).

One crucial aspect of RL is an individual’s ability to modulate their effective learning rate in accordance with the current environment. Various forms of environmental uncertainty, such as underlying volatility or reversal in the association between states or actions and outcome contingencies, require dynamic adjustments to learning by incorporating recent information more quickly into value expectations. Humans are remarkably adept at appropriately adjusting learning rate to different environments (10,11), but several studies have linked higher levels of anxiety or internalizing psychopathology shared across anxiety and depression to impoverished flexibility of learning (5,12,13).

One way in which effective learning rates may be modulated is by an adjustment of the relative recruitment of different neural processes within the brain. In particular, working memory (WM) processes are known to contribute to fast learning behavior in parallel to slower RL processes, and thus to modulate the effective apparent learning rate as a function of load (14–16). While RL processes have been neurally linked to several brain regions including the ventral tegmental area (VTA), the striatum and cortico-basal-ganglia circuitry (17,18), WM processes have primarily been linked to the brain’s prefrontal cortical executive control systems (19–22).

Although reinforcement learning has often been studied without consideration of WM, research has shown that reinforcement learning processes are supplemented by WM systems (14,16,23,24), and that WM processes may additionally interfere with RL (14,25,26). In particular, early-stage learning acquisition has been found to feature reduced activity of canonical RL neural systems such as cortico-basal-ganglia circuits and heightened activation of frontal cortical areas when compared with later-stage learning (16,17,27,28). Given that elevated activation in the prefrontal cortex (PFC) has consistently been found during (and may coordinate) the use of working memory (19–21), this may suggest that WM systems play a particularly key role in early-task RL learning acquisition.

To account for the parallel and contributory processes of working memory during reinforcement learning, a line of recent research has successfully introduced a new experimental paradigm which differentially manipulates the relative contributions of working memory and reinforcement learning systems during a learning and retention task. These studies have shown that jointly modeling the WM and RL processes can reveal key features of learning which cannot be accounted for with stand-alone RL models (15,29). Indeed, past research on individual differences has shown that using the RLWM framework can help more precisely identify the distinct mechanisms underlying apparently similar learning impairments; for example, learning impairments in schizophrenia appear driven by WM capacity limitations (15), in older adults by faster WM decay (30), and in young children by slower RL and weaker WM involvement (29). Thus, failure to take WM processes into account might also contribute to inconsistencies in findings within the anxiety RL literature.

The need to consider the contribution of working memory processes to anxiety-related deficits in learning is further supported by findings directly linking anxiety to aberrations in the use of aspects of prefrontal cortical executive systems (31,32) including specific deficits in working memory (33). In a study of N-back task performance in both safe and threatening environments, patients diagnosed with anxiety disorders (ADs) had impaired performance and reaction times across both environments (34). In addition, AD patients showed impaired recruitment of prefrontal cortical regions during these WM tasks. Other studies examining the same frontal regions have also hinted at a dimensional specificity to the role of anxiety in impaired executive function. An fMRI investigation of performance and cortical activity during a sustained attention task found that a general measure of trait anxiety was linked to impoverished recruitment of frontal regions at points where adjustments of attentional control were required, whereas a specific measure of worry separately impacted frontal-default mode connectivity during periods where attention lapsed (32). In another study, only anxious arousal, and not other indices of anxiety or negative affect, was linked to impaired perceptual decision-making (35). These findings highlight the possibility that concurrent investigation of different subdimensions of anxiety may be crucial for understanding anxiety-related differences in cognitive function.

Here we sought to use the enhanced RLWM experimental and modeling framework to disentangle pure RL from working memory contributions to differences in learning as a function of dimensions of anxiety. Given prior findings indicating that physiological and cognitive dimensions of anxiety may be differentially associated with impairments in processes spanning attentional control, working memory, and reinforcement learning, we sought to test our hypotheses with respect to specific characterizations of each symptom domain. We used scores on self-report measures of anxious arousal (Mood and Anxiety Symptom Questionnaire (36) (MASQ); Anxious Arousal (AA) subscale) and worry (Penn State Worry Questionnaire (37) (PSWQ)) to evaluate the influence of these two distinct dimensions of anxiety on learning and working memory involvement during reinforcement learning. Given the high rate of comorbidity between anxiety and depression, we performed a supplemental exploratory analysis of the relationship between the two anxiety measures mentioned, plus two common self-report measures of depressive affect (CES-D (38) and BDI-II (39,40), with all the parametric mechanisms of the winning RLWM model to inform potential directions for future research across the two highly comorbid disorders.

Finally, to highlight the importance of factoring in WM contributions to learning, we considered the effect of fitting RL models without WM modules to the experimental data for the current study. Here we examined two classes of RL models (using single or set size dependent learning rates) with varying parameterizations to include mechanisms of choice perseveration, forgetting, and negative feedback neglect. We assessed the mechanistic attribution of impaired learning performance and compared/contrasted it to the findings from the best-fit joint-process RLWM model to determine whether the RLWM model was able to improve specificity of findings relative to more common RL model formulations.

## Results

### Accuracy

In the current study, we employed a deterministic reward-based stimulus-action learning task which presents varying set sizes of stimuli by block, with stimulus set sizes (nS) ranging from 2 to 6 (see Figure 1a), in order to differentially engage WM versus RL systems during learning. Following a distractor task, a surprise test phase was presented which measured learning retention after WM had decayed. A total of n=164 participants were included in the analysis; see Methods for complete details.

**Figure 1.**
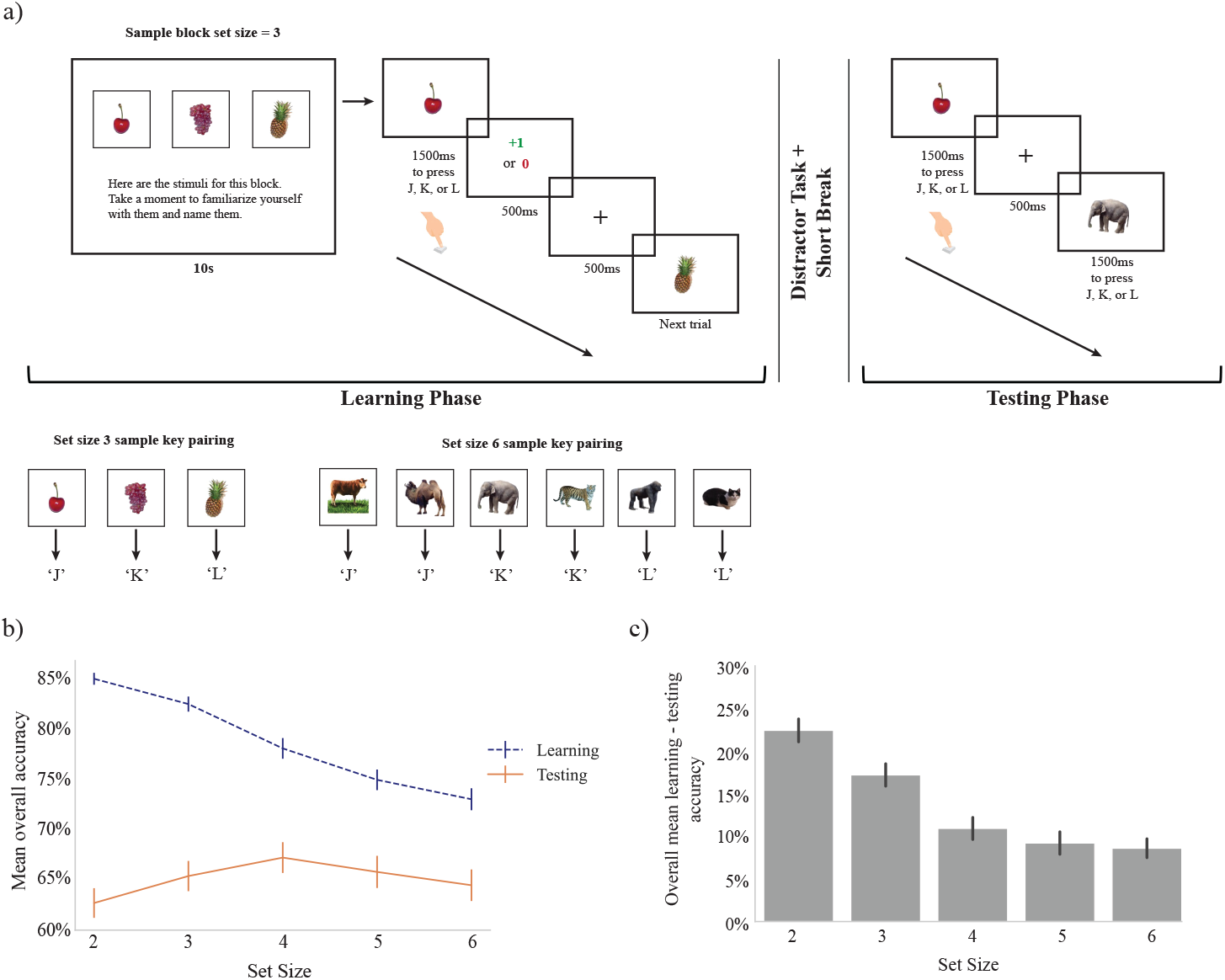
a) RLWM task. Participants completed 13 blocks of trials. Each block had a set size of 2, 3, 4, 5, or 6 images to be learned, and each stimulus was shown 13 times within each block for up to 1.5s each presentation (terminating when a key was pressed). Images for a block were related to each other (e.g., “Instruments”) and were unique for each block. Each image in a block was associated with one correct response key from 3 possible keys (‘j’,’k’,’l’). Participants learned the correct key to press for each image via trial and error, with immediate feedback displayed for 500ms after each key press (+1 if correct, 0 if incorrect). After a distractor task (an N-back task using unrelated shapes, not used for main analysis), participants were given a surprise testing phase in which each stimulus from the learning phase was shown 3 times (in shuffled random order across all stimuli). No feedback was provided in the testing phase. b) Mean overall accuracy during learning was significantly lower for high versus low set sizes, while mean overall accuracy during testing was not significantly modulated by set size. c) There was a significant drop in performance between mean overall learning versus testing accuracy at each set size; this difference was negatively associated with increasing set size.

We first sought to replicate previous findings of joint RL and WM involvement at a group level. During learning, participants performed better than chance (chance = 1/3), with mean accuracy across all set sizes of 78.2%. All set sizes had high mean overall performance above 70% accuracy (see Figure 1b), with set size 2 having the highest mean performance (mean=85%) and set size 6 the lowest (mean=75%). In line with previous literature, we observed that as set size increased, accuracy decreased: Overall set size slope (see Methods) median was 0.204 and was significantly greater than 0 for the group (Wilcoxon one-sample test statistic = 571.0, p = 0.000), indicating a significant effect of set size in (decreasing) performance accuracy.

The testing phase followed a distraction task, the aim of which was to eliminate the contribution of working memory to performance. In line with this, participant performance in the testing phase was substantially lower than during learning (Wilcoxon one-sided test statistic = 13003.0, p = 6.42e-25). Overall performance during testing still exceeded chance (chance = 1/3) with a mean accuracy of 65.2% across participants and set sizes. Despite greater accuracy during the learning phase for small set size stimuli, accuracy during testing was not significantly modulated by set size (set size slope median = −0.017, Wilcoxon one-sided test statistic=6176.000, p=0.334), reflecting a greater relative retention of learned associations at higher set sizes (see Figure 1b). Indeed, the difference in performance between learning and testing (which effectively factors out set size effects in learning accuracy) was significantly negatively associated with set size (Kruskal-Wallis = 80.075, p = 1.65e-16; see Figure 1c). This superficially counter-intuitive finding replicates previous findings and is consistent with greater reliance on working memory in the initial learning phase at smaller set sizes, but also of interference of WM blocking RL in smaller set sizes (14,30,41,42).

### Relationship of anxiety with performance

We used the MASQ AA subscale, which assesses self-report items specific to anxious arousal, to obtain a measure of physiological anxiety. The overall group mean score (+/−standard deviation) on the MASQ AA was 27.25 +/−10.86; see Supplements Figure 1). Scores on the MASQ AA subscale were significantly negatively correlated with mean overall performance during learning (Spearman rho(162) = −0.299, p = 9.79e-5; see Figure 2a for illustration using median split on MASQ AA scores). Further, individuals with higher MASQ AA scores also showed significantly greater effect of set size (as measured by set size slope; see Methods) on performance during learning (Spearman rho(162) = 0.220, p = 0.005). See Figure 2g for illustration of learning curves by set size using median split on MASQ AA scores.

**Figure 2.**
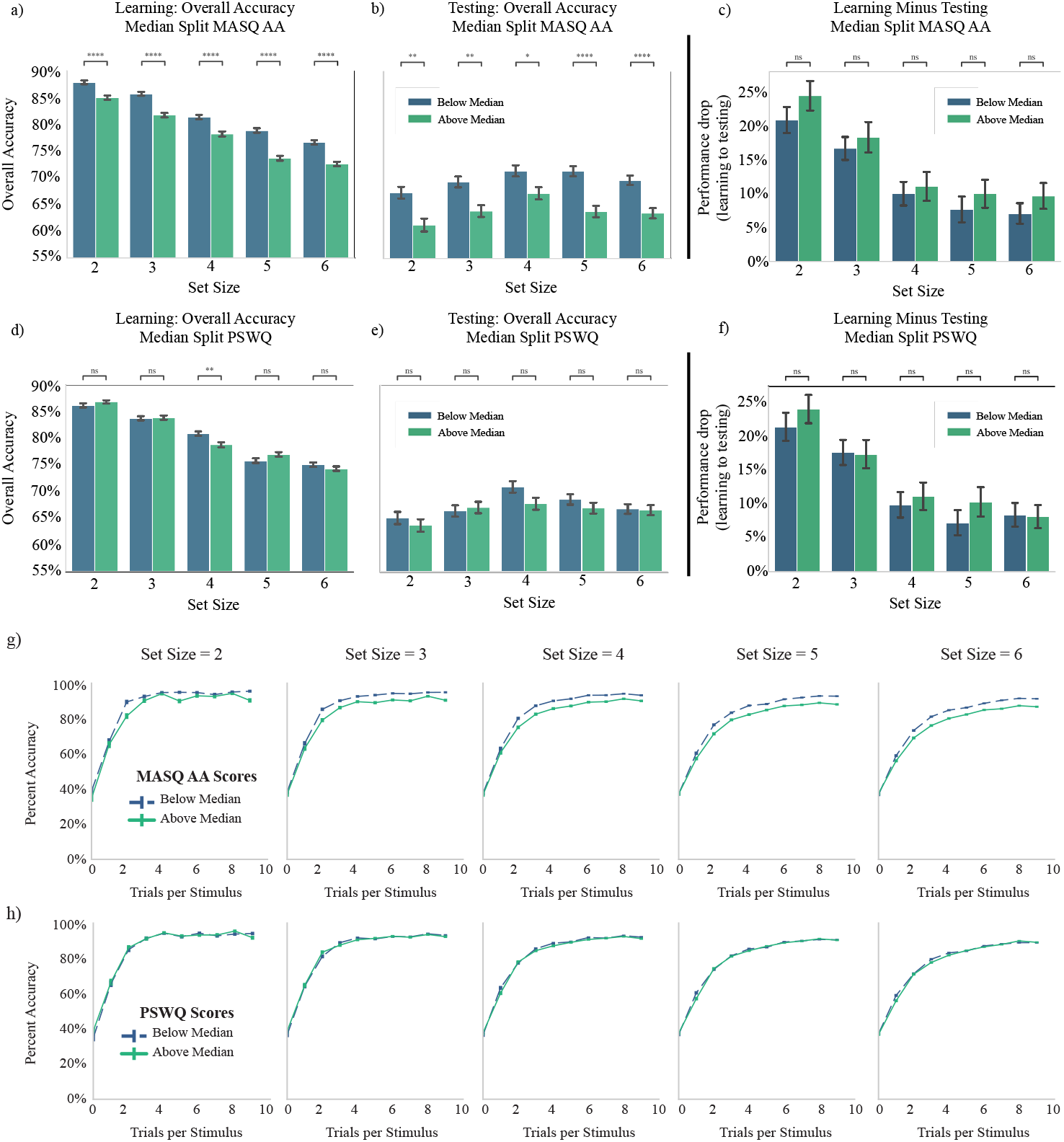
Median split of learning and testing performance on measures of trait anxiety. a,b) Individuals with above-median scores on the Mood and Anxiety Symptom Questionnaire (MASQ) Anxious Arousal (AA) subscale show significantly lower performance accuracy relative to below-median scores at every set size during both learning (a) and testing (b) phases of the RLWM task. c) There was no significant difference in performance drop between learning and testing phases for above-median MASQ AA relative to below-median MASQ AA scoring participants at any set size. d,e) Individuals with above-median scores on the Penn State Worry Questionnaire (PSWQ) do not have significantly different performance relative to below-median scores at any set size during learning except set size = 4 (but see Supplements Figure S.4.9), and no significantly different performance at any set size during testing. f) There is no significant difference in performance drop between learning and testing phases for above-median PSWQ relative to below-median PSWQ scoring participants at any set size. g) Learning curves for each set size split by median scores on MASQ AA reveal relative differences in learning speed and ultimate performance. h) Learning curves for each set size split by median scores on PSWQ reveal no relative differences in learning speed and ultimate performance.

As a complementary analysis, we conducted a repeated measures ANCOVA with within-subject factor of set size, covariate of z-scored MASQ AA scores, and dependent variable of mean learning performance. This revealed a significant within-subject effect of set size (within-subject set size effect with Greenhouse-Geisser correction applied F(3.430, 555.588) = 66.143, p < 0.001); a significant main effect of MASQ AA (between-subjects MASQ AA effect F(1,162) = 37.729, p <0.001); and a significant interaction of set size with MASQ AA (within-subject set size x MASQ AA effect with Greenhouse-Geisser correction applied F(3.430, 555.588) = 3.844, p = 0.007).

We next investigated whether physiological anxiety also affected long term retention in the testing phase. Indeed, higher scores on the MASQ AA subscale were also significantly associated with reduced overall testing performance (Spearman rho(162) = −0.289, p = 0.000; see Figure 2b for illustration using median split on MASQ AA scores). Of note, MASQ AA scores were not associated with any effect of set size on performance during the test phase (Spearman rho(162) = 0.099, p = 0.205). This may reflect the differing role of WM during learning and testing and hints at the possibility that high and low MASQ AA participants might show differential reliance on WM during learning. A repeated measures ANCOVA with within-subject factor of set size, covariate of z-scored MASQ AA scores, and dependent variable of mean testing performance revealed a significant main effect of set size (within-subjects set size effect with Greenhouse-Geisser correction F(3.744, 606.481) = 4.000, p = 0.004) and a significant main effect of MASQ AA scores (between-subjects MASQ AA effect F(1,162) = 17.956, p < 0.001), but only a weakly trending effect of interaction between set size and MASQ AA (within-subjects set size x MASQ AA effect with Greenhouse-Geisser correction F(3.744, 606.481) = 2.046, p = 0.091).

To more directly investigate whether the effect of physiological anxiety on testing performance was due to differences during learning, we next analyzed test phase accuracy relative to asymptotic learning phase accuracy. Higher MASQ AA subscale scores were significantly associated with reduced relative asymptotic performance between set size 2 and set size 6 during learning (Spearman rho(162)=0.207, p = 0.008), but not with relative overall performance between set size 2 and set size 6 during testing (Spearman rho(162) = 0.079, p = 0.314). We additionally tested the relationship between MASQ AA subscale scores and drop between asymptotic accuracy over last 3 trials during learning and overall performance during testing. A repeated measures ANCOVA with within-subject factor of set size, covariate of z-scored MASQ AA scores, and dependent variable of drop in performance between mean of final 3 learning trials and mean overall testing performance revealed a main effect of set size (within-subjects set size effect F(4,648) = 8.967, p < 0.001), but no main effect of MASQ AA (between-subjects MASQ AA F(1,162) = 0.135, p = 0.713) or interaction of MASQ AA with set size (within-subjects set size x MASQ AA effect F(4,648) = 0.537, p = 0.708). See Figure 2c for illustration using median split on MASQ AA scores. This may indicate that testing phase effects were primarily driven by differences in learning experience.

Unexpectedly, scores on the cognitive measure of anxiety (the PSWQ) did not appear to significantly impact task performance. The overall group mean score (+/−standard deviation) on the PSWQ was 52.12 +/−13.81; see Supplements Figure 1). During learning, PSWQ scores were not significantly associated with mean overall aggregate learning performance (Spearman rho(162) = −0.091, p = 0.248), but had a trending relationship with learning phase set size slope (Spearman rho(162) = 0.149, p = 0.057). There were no significant effects of PSWQ scores on performance during testing, either across set sizes (Spearman rho(162) = −0.074, p = 0.347) or as a function of set size (Spearman rho(162) = 0.009, p = 0.910). See Figure 2d-f, 2h for illustrations using median split on PSWQ scores. See Supplements for the associated ANCOVA results; here neither the main effect of PSWQ nor the interaction of PSWQ by set size was significant for either phase of the task. See Supplements for additional PSWQ analysis.

### Computational modeling

To investigate the computational mechanisms underlying learning and testing performance, we fitted a series of RLWM models (14–16) to the study data. Varying combinations of parameters were considered (see Methods), and model fit was measured by the Akaike Information Criteria (AIC; see Methods).

The model which best represented the overall behavioral data as measured by (lowest) total AIC was Model #5 (RLWM*_η*_*asym*_*_2ρK*; see Table 1, Figure 7, and Methods). This model included the following parameters: RL learning rate *α*_*RL*_, which applied when reward = 1 (RL learning rate = 0 when reward = 0 in winning model variant); stochastic choice noise *ε*; softmax inverse temperature for testing phase choice (*β*_*test*_; softmax inverse temperature for learning phase was fixed at *β*=50 to improve parameter fitting, and choice noise in learning is captured by *ε*); bias parameter *η*_*WM*_ which controlled negative feedback neglect in WM module; WM decay/forgetting parameter φ_*WM*_; and parameter *i*, which controlled information sharing between WM and RL for RL prediction error calculation. See Supplements Figure 2 for distributions of parameter values. Based on recent work regarding potential bias in some RL models (43), we performed a robustness check of the winning model by confirming that the addition of a perseveration choice kernel (Model #6) did not improve model fit, and did not result in significant perseveration or significant changes in main model parameters; see Supplemental Analysis and Supplements Figure 5 for details.

**Table 1.** Computational models of reinforcement learning evaluated. For each model: model key, list of parameters, short description, and total group AIC are shown.

| Model | Parameters | Description | Total AIC |
| --- | --- | --- | --- |
| Model #1:<br>RLWM | $\alpha, \beta_{\text{test}}, \varepsilon, \rho, \varphi, K, i$ | Basic RL and WM model with parameter $i$ to coordinate information sharing between modules | 146,311 |
| Model #2:<br>RLWM<sub>bias</sub> | $\alpha, \beta_{\text{test}}, \varepsilon, \rho, \varphi, K, \eta, i$ | Same as RLWM, but updates following negative feedback are partially neglected according to parameter $\eta$ | 143,910 |
| Model #3:<br>RLWM<sub>bias_2r</sub> | $\alpha, \beta_{\text{test}}, \varepsilon, \rho_{\text{low\_ss}}, \rho_{\text{high\_ss}}, \varphi, K, \eta, i$ | Same as RLWM<sub>bias</sub>, but allows WM choice weight to vary between set sizes below versus above capacity | 143,799 |
| Model #4:<br>RLWM<sub>asymbias</sub> | $\alpha, \beta_{\text{test}}, \varepsilon, \rho, \varphi, K, \eta_{\text{asym}}, i$ | Same as RLWM<sub>bias</sub>, but negative feedback is neglected asymmetrically for RL (totally neglected) versus WM (partially neglected via $\eta$) | 143,718 |
| Model #5:<br>RLWM<sub>asymbias_2r</sub> | $\alpha, \beta_{\text{test}}, \varepsilon, \rho_{\text{low\_ss}}, \rho_{\text{high\_ss}}, \varphi, K, \eta_{\text{asym}}, i$ | Same as RLWM<sub>asymbias</sub>, but allows WM choice weight to vary between set sizes below versus above capacity | 143,629 |
| Model #6:<br>RLWM<sub>asymbias_2r_ck</sub> | $\alpha, \beta_{\text{test}}, \varepsilon, \rho_{\text{low\_ss}}, \rho_{\text{high\_ss}}, \varphi, K, \eta_{\text{asym}}, i, \tau, \kappa$ | Same as RLWM<sub>asymbias_2r</sub>, but includes choice kernel to reflect decaying influence of past action choices on current choice | 143,770 |

The total AIC of the winning model (Model #5; 143,629) was very close to the total AIC of the second-best model (Model #4; 143,718). We therefore confirmed in supplemental analysis that all results below remained significant when tested in Model #4; see Supplemental Analysis.

### Model based analyses

We first verified that model parameters were identifiable in the winning model by simulating data with fixed parameter values and assessing the accuracy of the parameter values fit to the simulated data (see Supplements Figure 3). Additionally, we verified that models themselves were identifiable by performing a model recovery analysis, simulating three data sets for each of three selected models, and fitting each of the resulting nine data sets with each of the selected models to confirm that the generative model best fit its own data in each case (see Supplements Figure 4). Furthermore, model validation showed that the winning model (Model #5) could capture the learning dynamic and testing phase accuracy well (see Figure 3).

**Figure 3.**
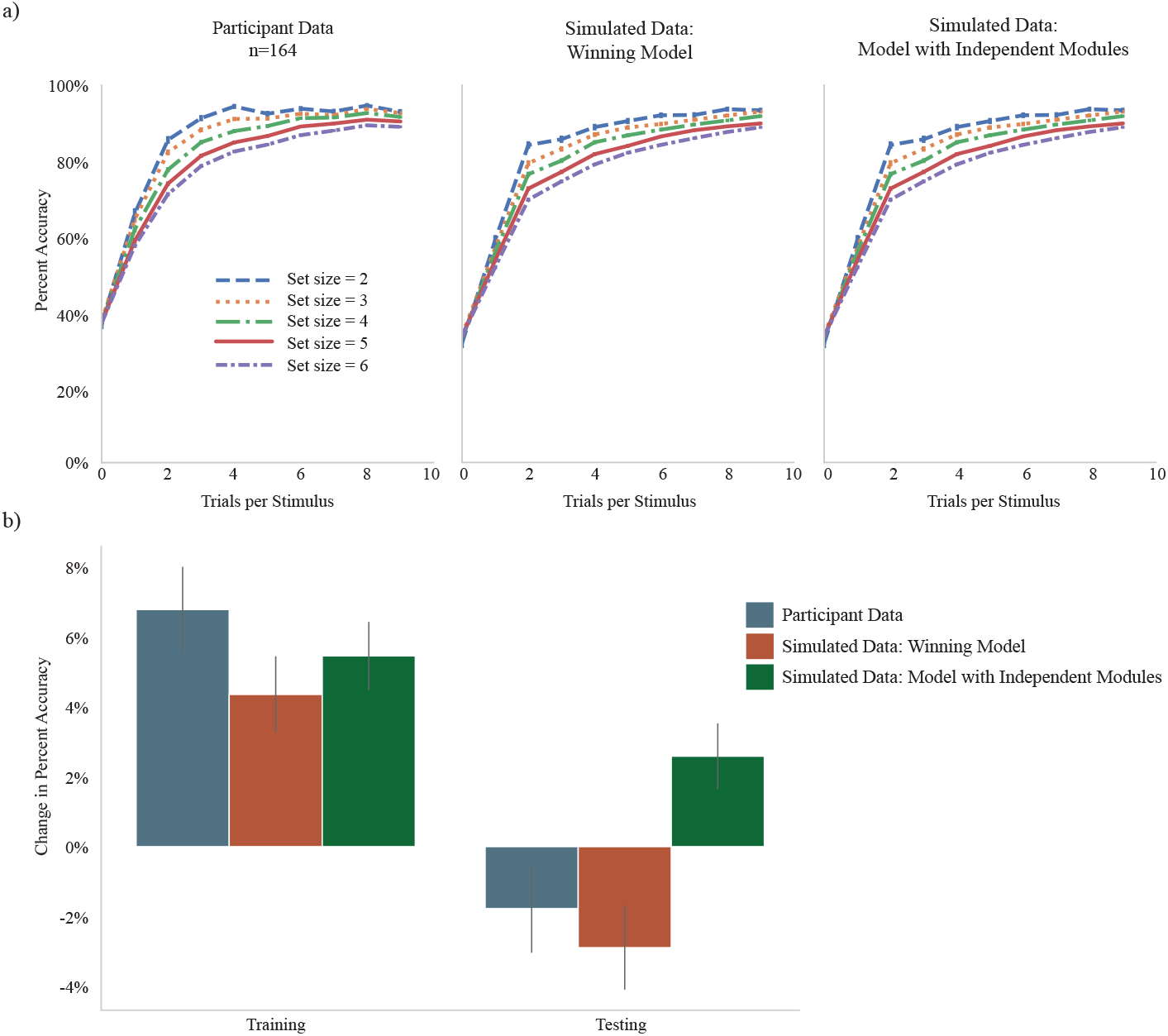
Model validation. a) Average learning curves across each set size represented as mean percent accuracy per stimulus presentation count. Left panel; actual participant data. Middle panel; data simulated using the winning model (Model #5). Right panel; data simulated using an RLWM model with fully independent RL and WM modules (for illustration). b) Performance difference between set size = 2 and set size = 6 (mean percent accuracy SS2 – mean percent accuracy SS6) in training (left) and testing (right) phases of the RLWM task. Participant data shows a reversal of this difference between training and testing. This pattern is reproduced by simulated data from the winning model (Model #5), but is not captured by model variants with fully independent RL and WM modules.

### Reduced learning and elevated WM decay associated with physiological anxiety

Given the significant behavioral impairment to learning and testing phase performance for higher scores on the MASQ AA, we examined the relationship of these scores with specific cognitive mechanisms of learning and working memory use as quantified by the winning RLWM computational model (Model #5). All results held when separately tested for significance in the second-best fitting model (Model #4); see Supplemental Analysis.

We performed initial hypothesis testing for participant scores on physiological arousal (MASQ AA scores) and cognitive worry/anxiety (PSWQ; for completeness) against one RL-specific parameter (learning rate *α*), two WM-specific parameters (forgetting φ_*WM*_, and neglect of negative feedback in WM *η*_*WM*_), and two parameters indexing relative contribution of WM to the policy (WM confidence at low set sizes *ρ*_*low*_, WM confidence at high set sizes *ρ*_*high*_) from the winning model (see Methods).

Higher MASQ AA subscale scores were significantly related to lower RL learning rate (rho(162) = −0.224, FWE-corrected p = 0.040) and higher rate of working memory decay (rho(162) = 0.223, FWE-corrected p = 0.040; see Figure 4). Prior to statistical correction for multiple comparison, higher MASQ AA subscale scores were also associated with a greater bias against the use of negative feedback in working memory, but this did not survive correction for FWE (rho(162) = 0.199, uncorrected p = 0.010, FWE-corrected p = 0.104). No significant associations were found between PSWQ scores and any of the specified parameter values.

**Figure 4.**
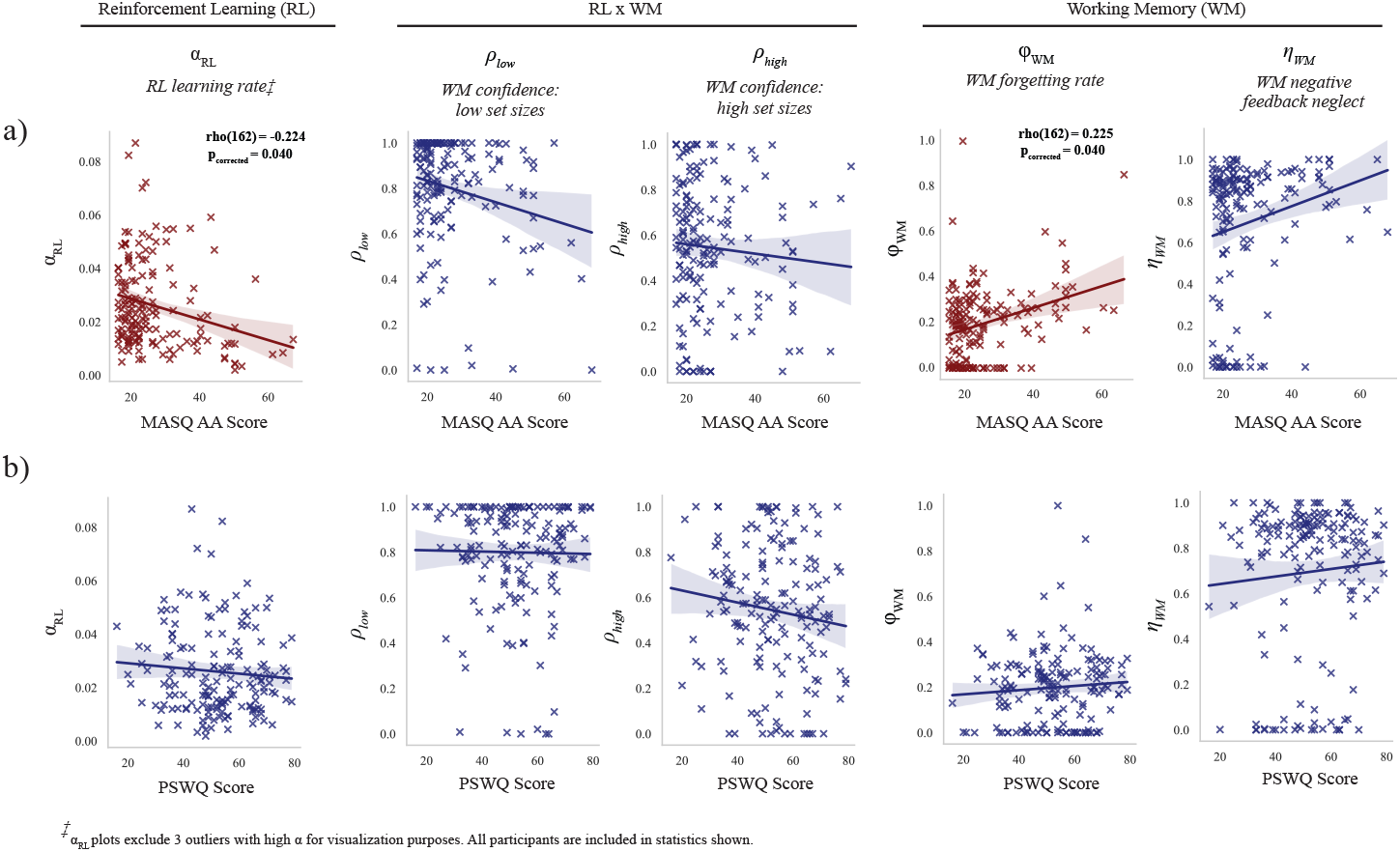
Relationship of learning and working memory model parameters (Model #5) with individual scores on MASQ AA (top row) and PSWQ (bottom row) From left to right, parameters tested: RL learning rate, WM forgetting rate, WM confidence weight for low and high set sizes, and WM bias against negative feedback updating.

### RL models without WM module vary in characterization of role of anxiety

Modeling learning with joint RL and WM processes revealed that two factors contributed to differences in learning as a function of MASQ AA. We performed an exploratory analysis investigating how our finding would be interpreted should modeling fail to account for WM. To do so, we additionally fit our data with a series of 10 variants of canonical RL-only models that did not include WM modules (see Methods). This allowed us to examine what our computational model-based findings would be if the data were fit with typical reinforcement learning models which do not account for WM. Given the structure of the task, which features 5 different set sizes across blocks, we considered simple RL models with either one shared learning rate across all set sizes (RL_*α* models) or with a separate learning rate for each set size (RL_5*α* models). In each class, we considered a base RL model (with parameters comprising learning rate(s), softmax inverse temperature for testing phase *β*_*test*_, and undirected choice noise *ε*), as well as four additional model variants which included various combinations of parameters for choice perseveration, reinforcement learning decay (forgetting), and negative feedback neglect; see Methods for details.

None of the RL-only models fit the data as well as the winning RLWM model; however, the model fit as measured by total (sum of) group AIC was in the same range as the RLWM class, with the best-fitting RL-only model fitting the data better (based on AIC comparison) than the simplest RLWM model (Model #1) (see Figure 5).

**Figure 5.**
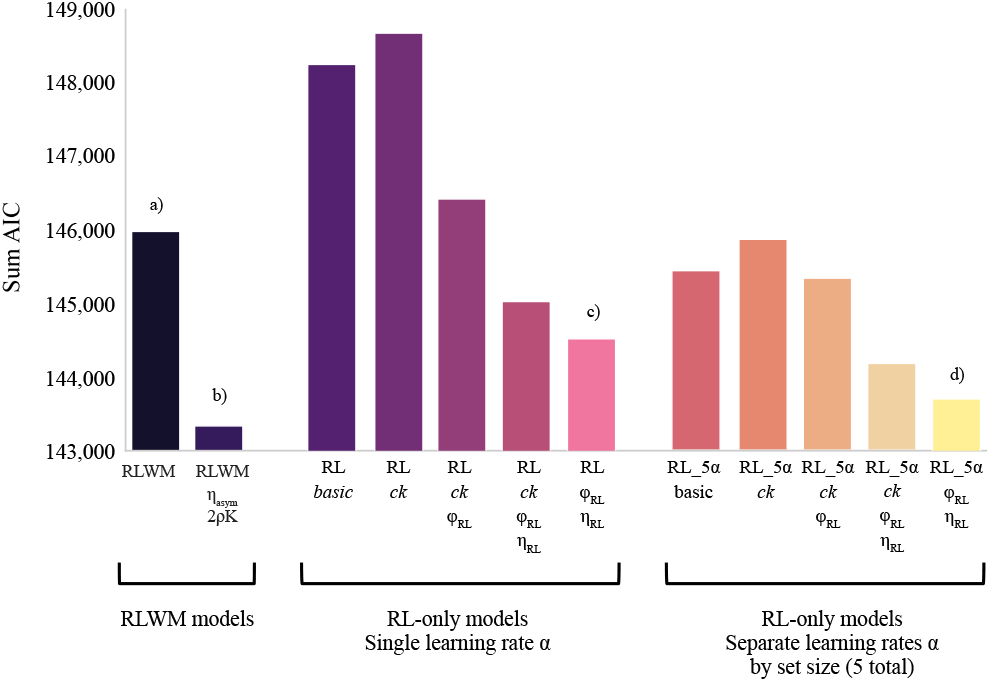
Model fit comparison (AIC) across base and winning RLWM model variants and 10 RL-only models tested. a) AIC for base RLWM variant. b) Winning RLWM model (Model #5) had best fit (lowest AIC) across all model variants tested. c) Best-fitting model variant in RL-*α* class. d) Best-fitting model variant in RL-5*α* class was best fit across all RL-only variants tested.

Within each class of RL-only models, the model variant which included forgetting (φ_*RL*_) and negative feedback neglect (*η*_*RL*_), but no choice kernel, fit the data best based on a comparison of AIC (see Figure 5). Across all RL-only models, the best-fitting model was the set size specific learning rate (5-alpha) version of this variant (RL_5*α_*φ_*RL*_*_η*_*RL*_, AIC = 144,170; see Figure 5).

Correlations between model parameters and scores on the MASQ AA varied somewhat across models depending upon specific parameterization; however, most models (8 out of 10) found a significant relationship between higher MASQ AA scores and higher undirected choice noise (*ε*) which was not present in the winning RLWM model, suggesting that RL-only models are likely to at least partially misattribute learning differences in anxiety to noise rather than deficits in learning or working memory (see Figure 6). The winning RL-only model (RL_5*α_*φ_*RL*_*_η*_*RL*_; see Figure 6) found that higher MASQ AA scores were significantly related to increased choice noise *ε*; rho(162)=0.238, p_corrected_=0.022), increased forgetting in RL φ_*RL*_; rho(162)=0.279, p_corrected_=0.003), and increased neglect of negative feedback (*η*_*RL*_; rho(162)=0.353, p_corrected_<0.001). See Figure 6 for significant parameter relationships with MASQ AA for each RL-only model as compared with the winning RLWM model (Model #5).

**Figure 6.**
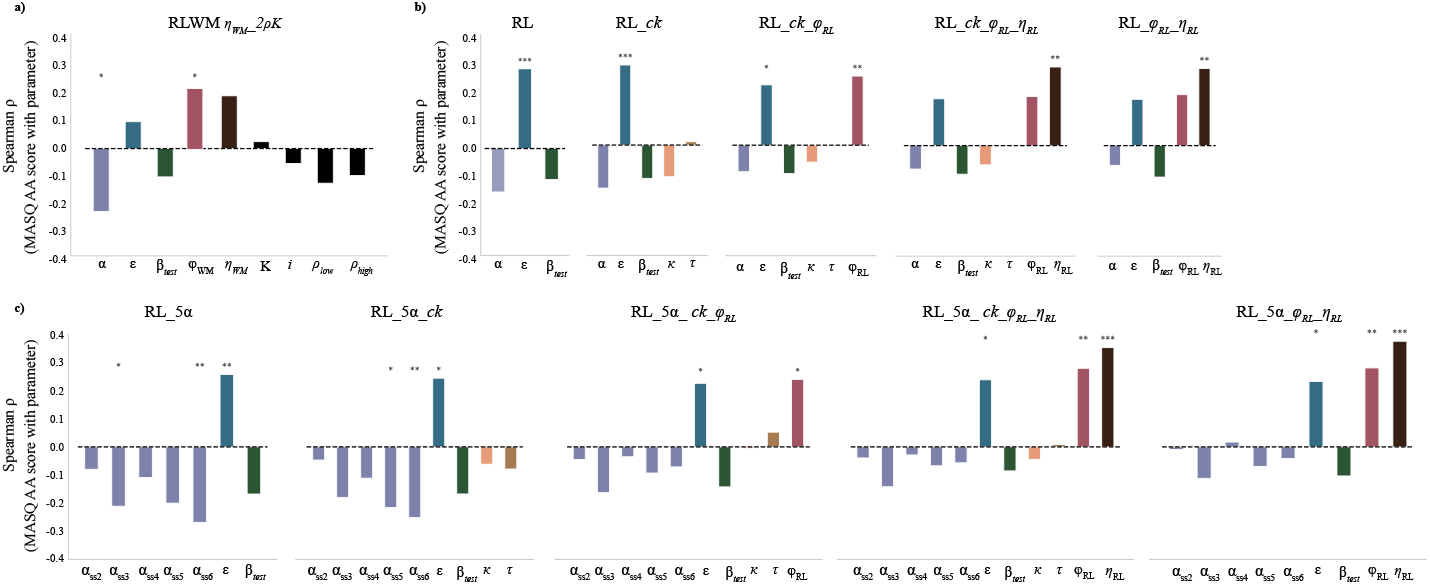
Comparison of relationship between MASQ AA scores and model parameters for winning RLWM model plus 10 RL-only model variants. Reinforcement learning models which do not specifically include the effect of working memory tend to attribute performance differences in high MASQ AA scoring participants to greater undirected choice noise, as well as to alternative mechanisms such as choice repetition and negative feedback neglect depending upon the specific model parameterization. a) Winning RLWM model from main analysis (Model #5) shows that higher MASQ AA scores are significantly related to reduced learning rate and increased working memory decay rate (forgetting). b) Results from 5 variants of RL-only models which feature a single learning rate parameter across all set sizes. Models considered include base model (learning rate (*α*), test phase inverse temperature (*β*_*test*_), and undirected choice noise (*ε*)), as well as incrementally including the following parameters: choice kernel (*ck*), RL forgetting (φ_*RL*_), and partial negative feedback neglect in RL (*η*_*RL*_). c) Results from 5 parallel variants of RL-only models which feature a separate learning rate for each set size.

**Figure 7.**
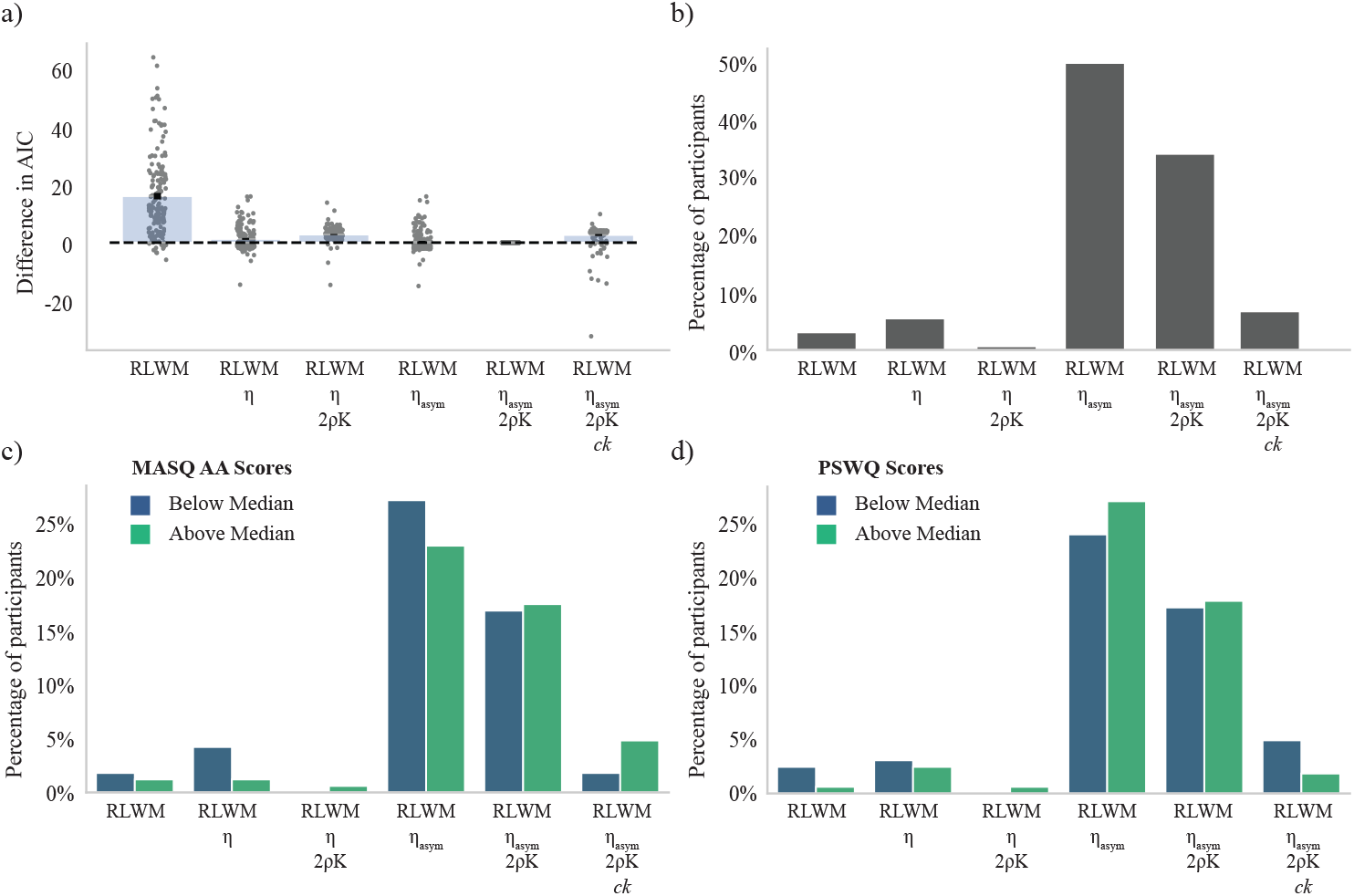
Model fit comparison using Akaike Information Criterion (AIC) a) Distribution of individual differences between model fit (AIC) for each model relative to individual AIC for the group winning model (Model #5). b) Percentage of overall participants best fit by each model. c) Percentage of participants best fit by each model by median split on MASQ AA subscale scores. d) Percentage of participants best fit by each model by median split on PSWQ scores.

## Discussion

Higher levels of anxiety have been separately associated with impairments in learning, working memory, and broader executive function in a number of studies (1,12,32,34,44,45). However, the influence of working memory on reinforcement learning processes in anxiety-related psychopathology has not to our knowledge been computationally investigated. In the current analysis, we used an experimental paradigm specifically designed to manipulate the relative load on learning versus working memory systems (16). We applied a computational model of reinforcement learning which accounts for the supportive role of working memory during learning to test whether relationships between anxiety and learning were attributable to learning processes, working memory processes, or both. We used two measures of anxiety-related psychopathology, one specifically characterizing physiological symptoms of anxiety and one characterizing cognitive symptoms of anxiety, to test any potential dimensional specificity in results.

Our findings indicate a complex picture of the relationship between anxiety, learning, and working memory. Firstly, we found no behavioral relationship between cognitive anxiety (as measured by PSWQ scores) and reduced performance in learning or testing across the task (with the exception of reduced learning at set size = 4). This was consistent with the results of our computational modeling analysis: no parameters of the winning RLWM model were found to have significant relationships with increased levels of cognitive anxiety.

In contrast, we found significant behavioral impairment in task performance at every stimulus set size across both learning and testing phases for individuals with higher levels of physiological anxiety (as measured by MASQ AA subscale scores). This performance impairment was mechanistically attributable to multiple parameterized mechanisms in computational modeling analysis. After correcting for multiple comparisons, MASQ AA subscale scores were related to a significantly lower learning rate and a significantly higher rate of working memory decay. This set of results suggests a multi-faceted relationship of anxiety with learning and its supportive processes, whereby both the learning process itself as well as the role of the supporting working memory systems are compromised in high levels of physiological anxiety.

Most studies of reinforcement learning within computational psychiatry have used paradigms and models which do not specifically account for working memory systems; indeed, our ability to model working memory contributions here is due to the experimental load manipulation, and is thus not easily applied retrospectively to existing reinforcement learning datasets. We therefore sought to assess what our findings for the current dataset would be when modeled without the inclusion of the working memory module. For comparison, we fit a series of more typical, RL-only models which did not account for contributions of WM to the dataset and examined the correlation between model parameters and MASQ AA scores for these model classes. Of note, 5 out of 6 RL-only models which included a parameter for forgetting (φ_*RL*_) found a significant association between rate of forgetting and MASQ AA scores, in parallel to the main analysis (Model #5) finding that higher MASQ AA scores were tied to faster forgetting in working memory processes (φ_*WM*_) (see Figure 6). Crucially, models which did not account computationally for WM contributions largely attributed the decreased learning performance for high MASQ AA scores to undirected choice noise (*ε*; 9 out of 10 models) and increased negative feedback neglect (*η*_*RL*_; 4 out of 4 models which included this parameter). The specific characterization of impaired mechanisms in higher MASQ AA scores varied depending on the particular parameterization of the models. Only 2 out of 10 models found a significant relationship between higher MASQ AA scores and lower learning rates, and in these models the relationship was only consistently observed at the largest set size (Figure 6).

These RL-only model analyses do not provide an exhaustive nor necessarily direct comparison with the RLWM models considered. In particular, we note that the winning RLWM model reflected total negative feedback neglect for RL and parameterized negative feedback neglect for WM, making it different from any possible characterization of RL-only models which can only include a negative feedback parameter for RL (their only module). We also note that this analysis may not be directly comparable to other RL-only studies in the field, both given the structural difference of the current task’s multiple set sizes across blocks when compared with typical static set size RL tasks, and in light of the purely deterministic (accurate) feedback used in the current study when compared with the (often) stochastic feedback structure of many RL tasks (46,47). Nonetheless, the comparison of attribution of anxiety-related learning task deficits between RLWM and RL models as applied to the current dataset points to the possibility that failure to model WM explicitly might lead to misleading interpretation of anxiety-related deficits in learning. An important area for future study lies in the investigation of potential interactions between individual differences in the effectiveness of working memory system contributions to learning and the specific demands placed on working memory by RL tasks of varying complexity and structure. This would valuably be extended to work with other clinical populations; for instance, individuals with ADHD are known to have anxiety levels well above the population average, and responses to medication in reversal learning performance have been shown to relate to WM capacity (48). Extension of the work conducted here might help to illuminate the mechanisms underlying such findings.

It will also be important to conduct similar investigations for other dimensions of internalizing psychopathology. In the current study, supplementary exploratory analyses of self-report measures of depressive affect (CES-D and BDI-II) revealed no significant relationships between these measures and any of the computational model parameters. We include these findings for completeness but note that a larger scale study is required to provide a fully powered investigation of additional dimensions of psychopathology, including but not limited to depressive affect.

In summary, the study findings provide insight into the potentially complicated relationship of anxiety with interactive systems of WM and learning. By revealing the emergent learning rate effects of physiological anxiety when working memory is jointly modeled with the learning process, and pointing to the reduced effectiveness of working memory’s contributions to learning due to quicker rates of forgetting in individuals with high levels of physiological anxiety, the current findings provides new insights into the problems anxious individuals have with learning while also highlighting the need, across the computational RL literature, to explicitly model cognitive processes such as, but not necessarily limited to, working memory when interpreting the behavior of healthy participants or investigating psychopathology-related alterations in task performance.

## Methods

### Participants

The study was conducted online with participants recruited via the UC Berkeley Research Participation Pool (RPP), which offers partial course credit to undergraduate students for participation in human subjects research. All participants completed online informed consent prior to participation. The study was approved by the UC Berkeley Committee for the Protection of Human Subjects (CPHS). An initial total of n=229 students [143=female and 86=male; mean age = 21.2 +/−2.4] participated prior to exclusions.

### Exclusions

Participants were eligible to participate in the online study if they confirmed that they were not currently taking antidepressant or anxiolytic medications and had not used cannabis within the preceding two weeks. Two attention checks were embedded in the self-report questionnaires (e.g., “Select 2 here to show that you are paying attention”) as data validity checks. Additionally, following Meade & Craig, 2012, once participants had completed the task they were informed that their participation credit was now guaranteed, and were asked to answer two questions honestly to ensure that the research would only use credible data: participants were asked to again confirm whether they had used cannabis in the last two weeks, and were asked whether they felt they had paid sufficient attention during the task that their data should be used in our study. 45 participants were excluded based on responses to the end-of-task questions, and 6 additional participants were excluded for missing two or more attention checks during the questionnaires.

Participants were also excluded if they had greater than two standard deviations 2SD) above the mean number of trial timeouts (>11.9% of trials; 6 participants excluded). Additionally, we calculated mean set size = 2 asymptotic performance across the last 5 same-stimulus presentations for set size = 2 blocks; participants with mean accuracy lower than 2SD below the group mean (<64% accuracy; 8 participants) were excluded from further analysis. Pre-exclusion population distributions for number of timeouts and asymptotic performance on set size = 2 were highly skewed (see Supplements Figure 7). Recent work exploring the effect of exclusions in highly skewed data on individual differences analyses has cautioned that overly aggressive exclusion criteria may introduce “shadow biases” into individual differences work by excluding participants disproportionately with respect to metrics of interest (50). Simulation work examining various exclusion criteria approaches for reaction time distributions, which are also highly skewed, showed that a 2SD cutoff was among the least biased methods for tail exclusions in these distributions (51).

The number of participants included in the final analysis was n=164 (104 = female; mean age 21.20 +/−2.43). 34 participants self-reported Hispanic or Latino ethnicity. Self-identified race distribution of participants was as follows: 94 Asian, 8 Black or African American, 34 White, 2 Native American or Pacific Islander, 12 who identified as “More than one race” and 14 who identified as “Unknown” race.

On a within-participant level, trials that timed out were excluded (mean number of timeout trials = 4.74), and subsequently any trial blocks for which a participant had fewer than 9 presentations of any stimulus remaining in a block after exclusions were also excluded (1 block excluded for each of 5 participants).

### Experimental paradigm

The behavioral task used was a variant of the classic RLWM task (Collins & Frank 2012). The main task comprised a learning (or “training”) phase, followed by an unrelated distractor task, and finally a surprise testing phase of the original task stimuli. During the learning phase, participants were presented with a series of stimuli (images) on screen, with one stimulus shown for each trial. There were three possible correct key presses associated with each stimulus; keyboard keys ‘j’, ‘k’, or ‘l’. Each stimulus within a block was associated with only one correct key press (but multiple stimuli within a block could be associated with the same correct key press). Participants had 1.5s to select an action for each stimulus presented; if no response was selected in the allowed time, a message “You did not make a selection in time!” was displayed for 500ms, and the task advanced to the next trial. If participants responded with a key press, the stimulus was removed from the screen and they were given accurate feedback (a green +1 for correct responses, or a red 0 for incorrect responses) presented for 500ms. A fixation cross was shown for 500ms between each learning trial with a 100ms blank screen immediately before and after the fixation cross. See Figure 1.

The learning task consisted of 13 blocks of trials. Within each block, participants learned the correct key press action for each stimulus in the block through multiple trials and feedback. Each trial block included a stimulus set size (nS) varying between 2, 3, 4, 5, or 6 stimuli. Stimuli for each block were drawn from a randomly selected image category (such as ‘nature’, ‘shapes’, ‘musical instruments’, etc.) without replacement, such that each set of unique stimuli within a block were of the same category and no stimuli or categories were repeated across blocks.

Participants were randomly assigned to one of 10 pre-generated learning task trial sequences. In each generated trial sequence, participants started and ended with a block of set size = 2 to mitigate the possible conflation of primacy and recency effects with working memory. Block 2 was always of set size = 3; the remaining intermediate blocks were shuffled so that each set size was presented once in blocks 3-7 and once in blocks 8-12. Each stimulus for a given block was shown 13 times within that block, so the total number of trials per block varied with stimulus set size. Stimuli sequences within each block were generated pseudo-randomly controlling for a uniform distribution of delay between each two successive presentations of the same stimulus across every [1:2*nS] trials per block.

Following the learning phase, participants were given an optional break of up to one minute, followed by a distractor task. The distractor task used in the current experiment was a short n-back task in which participants were shown a series of images (shapes which were not used in the learning task) and asked to press the left arrow key of the keyboard if the image on screen was the same as the image that had appeared “n” stimuli ago. Participants completed a 1-back, 2-back, and 3-back task (mean completion time = 18.8 minutes). Following the n-back task, participants were given an optional 30 second break.

Following this break, participants were informed that they would be tested on their learning of the actions associated with the stimuli they had seen during the earlier learning portion of the experiment. Participants were shown each stimulus from their learning phase a total of three times during the testing phase. In each testing trial, the stimulus appeared on screen for 1.5s, and the participant selected ‘j’, ‘k’, or ‘l’ based on their earlier learning for the stimulus. Importantly, participants were not given feedback on accuracy in the testing phase, so additional learning could not occur. If participants did not select an action quickly enough, a message “You did not make a selection in time!” was displayed for 500ms. A fixation cross was shown for 500ms between each testing trial with a 100ms blank screen immediately before and after the fixation cross.

### Self-report measures of trait anxiety and depression

Prior to the behavioral task and following informed consent, participants were asked to complete a short series of questionnaires designed to measure levels of anxious and depressed symptomatology. The questionnaires used in this study comprised the following: the State-Trait Anxiety Inventory trait subscale (STAI-T; Spielberger, C. D., Gorsuch, R. L., Lushene, R., Vagg, P. R., & Jacobs, G. A., 1983); the Penn State Worry Questionnaire (PSWQ; Meyer et al., 1990); the Beck Depression Inventory (BDI-II; Beck, 1961); and the Mood and Anxiety Symptom Questionnaire anxious arousal subscale (MASQ AA; Watson & Clark, 1991). Items addressing suicidality were excluded. Final number of items included was n=126.

Scores on the MASQ AA subscale had a mean of 27.25 (standard deviation=10.86), with a minimum participant score of 17 (minimum possible scale score=17) and a maximum participant score of 68 (maximum possible scale score=85). Scores on the worry measure PSWQ had a mean of 52.12 (standard deviation =13.81), with a minimum participant score of 16 (minimum possible scale score = 16) and a maximum participant score of 79 (maximum possible scale score = 80). These subscale scores differentiate between anxiety as characterized by somatic symptoms versus cognitive symptoms and were used in primary hypothesis testing (see Results). Complete score distributions for the participant group for each questionnaire measure, including the BDI and STAI-T scales used in exploratory post-hoc analyses, are shown in Supplemental Figure S1.

### Behavioral analysis

Previous investigations of learning under various set sizes have found a robust and highly replicable effect of set size on learning over time (14,16,29). To confirm that our data replicated these established results, we first examined the mean learning curves across participants at each set size, measured as mean percent response accuracy by stimulus presentation. Additionally, we calculated overall learning accuracy as the mean accuracy across all presentations of all stimuli for each set size, and asymptotic learning accuracy as the mean of the last 5 presentations of each stimulus across blocks with the same set size. Overall testing accuracy was calculated as the mean accuracy across all stimuli presentations (each image was shown 3 times during testing) for each set size.

To quantify the influence of increases in set size on participant performance, we calculated a set size slope for each participant (following (29) according to the following equation:

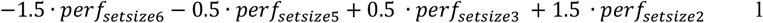

Where underlying data followed a normal distribution, we used Pearson correlations and t-tests as specified throughout the text. We used Spearman rank correlation, Wilcoxon, Kruskal-Wallis and Mann-Whitney U testing as specified throughout the text to test various correlations and distributional differences. Statistical tests were performed in Python and R.

### Computational modeling

We modeled learning and testing performance using a series of dual-module reinforcement learning and working memory (RLWM) models of varying levels of complexity. This class of models has been shown to effectively tease apart the effects of working memory from those of reinforcement learning in RLWM tasks with a set size manipulation as described above (14–16,29). Model variants were tested individually and in combination; see Model key for details.

#### Baseline model: Reinforcement learning and working memory (RLWM)

Our baseline model was a two-module reinforcement learning (RL) and working memory (WM) model in which separate RL and WM processes contribute collaboratively to learning and decision-making (14,42). We note that while some previous studies investigating learning-phase data from the RLWM task have used models in which the RL and WM modules operate independently (e.g., (16), these models are not capable of capturing participant testing phase performance (see Figure 4 of the current paper) and so are not considered here.

The WM module tracks weights for each possible action (*a*) given each stimulus (*s*) per block. Working memory weights (denoted as *W*(*s, a*)) are initialized to random chance (1/*nA*, where *nA=3* is the number of possible actions per state) and are updated after each trial based on feedback assuming perfect information retention. Although working memory has high information retention in the short term, the stored weights are assumed to decay rapidly at a parameterized rate of φ between each trial update in order to reflect the short-term nature of WM.

The weight for the current action/stimulus *(s*_*t*_, *a*_*t*_) for each trial is updated based on a prediction error (equation 1 below) from the observed trial feedback *r*^*t*^ (1 = correct, 0 = incorrect) with complete retention (learning rate *α*_*WM*_ = 1) such that:

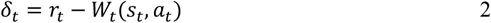

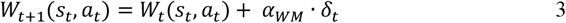

WM weights for all stimuli and actions *(s*,*a)* decay back toward initial values after each trial according to:

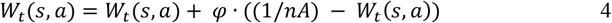

The contribution of working memory to action selection in each block is weighted according to a prior WM confidence weight parameter *ρ* ∈ [0,1] and WM capacity parameter *K* ∈ [2,6] compared to the number of stimuli in the set size (ns) for the current block.

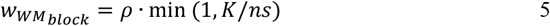

Meanwhile, the reinforcement learning (RL) module tracks action values (denoted *Q*_*RL*_(*s, a*)), also initialized to 1/*nA*. Values are updated for the current action/stimulus at each trial using a cooperative approach whereby both RL values and WM weights contribute, with a reward prediction error (*rpe*) defined as follows:

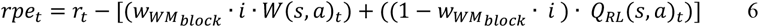

and parameter *i* ∈ [0,1] controls the strength of the information sharing between RL and WM modules, such that WM knowledge contributes to the expectation portion of the prediction error.

Values for RL are then updated using a learning rate parameter *α*_*RL*_ ∈ [0,1] governing the rate at which feedback is incorporated into the estimate.

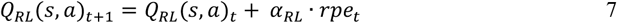

To model the probability that an agent selects a given action at a given time, we use a weighted mixture softmax choice policy to evaluate the relative value of each action during the decision process. The probability of selecting action *a* following stimulus *s* is denoted as follows:

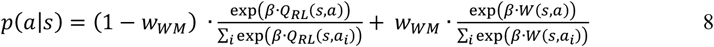

where the parameter *β* ∈ [0,100], referred to as the softmax inverse temperature, controls the extent to which relative action values (as opposed to stochastic choice noise) are used in decision choice. During learning, the softmax inverse temperature was set to a fixed value of *β* = 50 to improve parameter reliability (14) and choice noise was captured via an undirected noise parameter *ε* ∈ [0,1], such that:

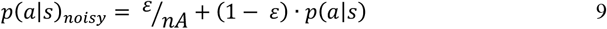

Modeling of choice selection during the testing phase used the final learning phase *Q*_*RL*_ values for the test stimuli, and assumed an epsilon-noisy softmax action selection with no contribution of working memory to choice. The parameter *β*_*test*_ ∈ [0,100] was fit in the testing phase softmax action selection policy (reflecting potential weakening of RL-values’ influence on test), while *ε* was fit jointly between the learning and testing phases (reflecting a shared tendency for lapses or inattention).

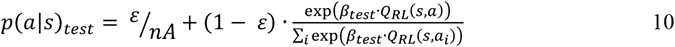

#### Negative feedback neglect variant (“_*bias*”)

Previous studies (14) have found that individuals often update action values more slowly following negative feedback than following positive feedback during learning. In *_bias* variant models, a parameter *η* ∈ [0,1] is introduced which reduces the learning rate for both RL and WM modules following negative feedback.

We parameterize this by reducing the learning rate in both RL and WM modules following negative feedback according to the rule:

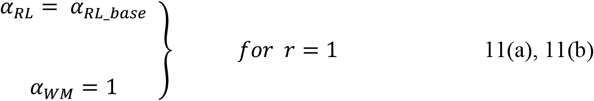

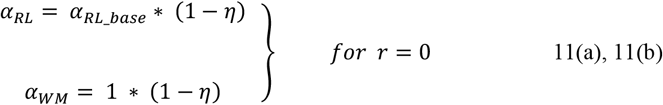

#### Asymmetric negative feedback neglect variant (“_*asymbias*”)

In the asymmetric negative feedback neglect model variant, we tested the hypothesis that individuals entirely neglect negative feedback in the RL module while still allowing for biased neglect in the WM module.

We parameterize this by reducing the learning rate asymmetrically for RL and WM modules following negative feedback according to the rule:

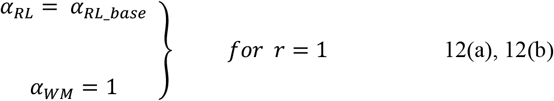

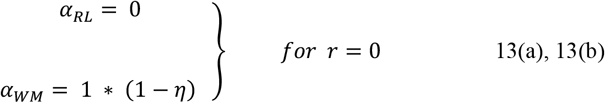

#### Split WM confidence variant (“_*2r”*)

In the split WM confidence model variant, we expanded the WM prior confidence parameter *ρ* into two parameters, *ρ*_*above_K*_ and *ρ*_*below_K*_. This modification allowed the weight placed on working memory during choice selection to vary based on whether the current block set size exceeded an individual’s working memory capacity *K*.

The working memory weights for this model variant were then calculated according to:

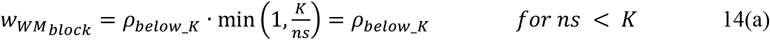

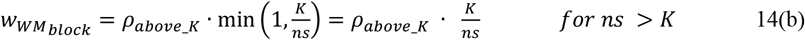

#### Choice kernel variant (“_ck”)

When making choices in learning tasks, participants may have a propensity to repeat previous actions regardless of value (a phenomenon sometimes referred to as perseveration or sticky choice). A choice kernel allows for multiple previous motor actions to influence the current choice, with a decaying influence of choices that are further in the past. In some (but not all) data sets, models which do not include action perseveration over multiple previous actions may induce a bias in parameter fitting, allocating variance inappropriately to asymmetric value updating following negative feedback (43,53).

We therefore tested a model variant which included the influence of past actions on current action choice during the learning phase of the task. The choice kernel (*Ck*) acts as a weighted trace of previous actions (i.e., pressing ‘J’,’K’, or ‘L’), and is initialized to 0 for each action.

Following each trial, the choice kernel is updated for the selected action and the stored action values decay at rate *τ* according to:

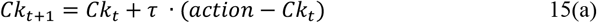

where *action* is an array of length 3 with value=1 for the chosen action and value=0 for each of the two unchosen actions on the current trial.

Each action decision is directly influenced by the choice kernel according to perseveration parameter κ:

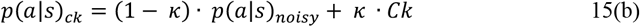

### Model fitting procedure

Model parameters were fit using maximum likelihood estimation in Matlab with the function fmincon to minimize negative log likelihood. Individual maximum likelihood estimates were performed using 20 independent randomly selected starting points to improve selection of global maxima. All parameters were constrained to [0,1], except the softmax inverse temperature for the testing phase, which was scaled to be constrained on the interval [0,100], and the working memory capacity parameter K, which was constrained to the continuous interval [2,6].

### Model comparison

We evaluated the fit of variants of the RLWM model with combinations of each modification described above. The final model space included 5 models as shown in Table 4.1.

Model fit was assessed by comparing the Akaike information criterion (AIC; Akaike, 1974) for each model in aggregate. The AIC measures overall model fit while penalizing for complexity, and we verify that it supports adequate model identification within the RLWM framework (29). The winning model with the lowest total AIC was Model #5 (RLWM*_asymbias_2r)*, with the following 9 parameters: RL learning rate, test phase softmax inverse temperature, epsilon undirected choice noise, WM confidence for low set sizes, WM confidence for high set sizes, WM forgetting, WM neglect of negative feedback, WM capacity, and an interaction parameter for cooperative updating between RL and WM systems.

There were substantial individual differences in best fit model, with Model #4 best-fitting a greater percentage of participants based on individual AIC (see Figure 4.2). Model #4 was nested within the model with the overall lowest AIC (Model #5), indicating that the additional parameters in Model #5 were not necessary to fit some participants but were necessary to best fit others. Of note is that, because some models are nested, the more complex model could yield the exact same maximum likelihood as the simpler model for individuals who did not require the additional parameters by finding null-effect parameter values as optima for those individual parameters, so any improvement in AIC between Models #4 and #5 for a given individual would be due only to the penalty for modeling more parameters than necessary for that individual in the more complex model.

We compared individual differences in model best fit for high versus low anxiety participants using a median split on each of PSWQ and MASQ AA score, which illustrated that individual differences across model fit were qualitatively related to scores on metrics of anxiety (see Figure 4.2). Given that Model #5 showed the overall lowest AIC, best fit a subset of individuals, and contained Model #4 as a nested component, we proceeded with Model #5 as the winning model for purposes of hypothesis testing. We performed post-hoc confirmatory analysis of the significant findings in the analogous parameters in Model #4 to verify that results were robust across the two models.

### Model validation

Model validation was performed by simulating data from a generative version of the winning computational model (Model #5, above). Learning curves by set size, difference in performance between low and high set size during learning, and difference in performance between low and high set size during testing were replicated by the model (see Figure 4).

Parameter recovery was performed for the winning model to test the identifiability of all model parameters. Participant data was simulated using the fit parameter values from the final model, and this simulated data with known underlying parameters was then fit using the optimization process described above. Recovered (fit) parameter values were then compared with the generative (known) parameter values. Parameters generally recovered well, with both significant parameters (learning rate and working memory decay) having correlations between generative and fit parameters greater than or equal to 0.80. See Supplements Figure S3 for parameter recovery analysis for each model parameter.

Model identifiability was confirmed via a model-recovery analysis. Three sets of participant data were simulated from each of the following models: RLWM_bias (Model #2), RLWM_asymbias (Model #4), and RLWM_asymbias_2r (winning Model #5). Each simulated data set was then fit by each of the three selected models, and the resulting best-fit model was compared to the generative model for that data. Models accurately recovered their generated data; see Supplements Figure S4 for model recovery analysis.

### Model-based hypothesis testing (main analysis)

We performed initial hypothesis testing for dimensions of anxiety characterized by physiological arousal (MASQ AA scores) and cognitive anxiety/worry (PSWQ). We tested participant scores on each measure against one RL-specific parameter (learning rate) and four WM-related parameters (forgetting, WM confidence at low set sizes, WM confidence at high set sizes, and neglect of negative feedback in WM) from the winning model. We performed nonparametric correlation analyses due to the non-normality of the underlying data. We report p-values corrected using the Bonferroni statistical correction for 10 simultaneous comparisons (2 trait scores across 5 parameters each) to account for potential family-wise error across multiple tests (and additionally report uncorrected p-values for comparison where specified).

### Additional modeling analysis (RL-only models)

For comparison and illustration (see Results), we fit 8 additional RL-only models which did not include WM modules to our data. Since the structure of our task included 5 different set sizes of stimuli across blocks, we considered one class of RL-only models with a single learning rate across set sizes (RL_*α* models) and one class of RL-only models with a learning rate for each set size (RL_5*α* models).

Four RL-only models were fit for each class; a base variant, and 3 additional variants which incrementally (cumulatively) added parameters describing the following mechanisms: stickiness of choice (*s*), RL forgetting (φ_*RL*_), and negative feedback neglect (*η*_*RL*_), formulated as outlined in the equations below.

The basic formulation of the RL-only models was directly analogous to the basic RLWM model (Model #1) with the WM components removed. For the RL_*α* base model, the reinforcement learning (RL) module tracks action values (denoted *Q*_*RL*_(*s, a*)), initialized to 1/*nA*. Values are updated for the current action/stimulus at each trial according to a reward prediction error (*rpe*) defined as:

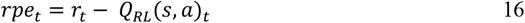

Values are then updated using a learning rate parameter *α*_*RL*_ ∈ [0,1] governing the rate at which feedback is incorporated into the estimate.

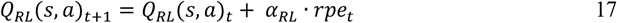

To model the probability that an agent selects a given action at a given time, we use a softmax choice policy to evaluate the relative value of each action during the decision process. The probability of selecting action *a* following stimulus *s* is denoted as follows:

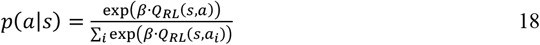

where the parameter *β* ∈ [0,100], referred to as the softmax inverse temperature, controls the extent to which relative action values (as opposed to stochastic choice noise) are used in decision choice. During learning, the softmax inverse temperature was set to a fixed value of *β* = 50 to improve parameter reliability (14) and choice noise was captured via an undirected noise parameter *ε* ∈ [0,1], such that:

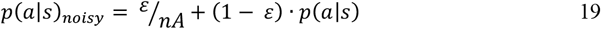

Modeling of choice selection during the testing phase used the final learning phase *Q*_*RL*_ values for the test stimuli and assumed an epsilon-noisy softmax action selection. The parameter *β*_*test*_ ∈ [0,100] was fit in the testing phase softmax action selection policy (reflecting potential weakening of RL-values’ influence on test), while *ε* was fit jointly between the learning and testing phases (reflecting a shared tendency for lapses or inattention).

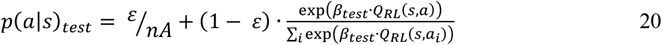

The RL_5*α* base model followed the same equations as the RL_ *α* base model shown above, but included 5 separate learning rate parameters *α*_*ss*2,_*α*_*ss*3,_*α*_*ss*4,_*α*_*ss*5,_*α*_*ss*6_ ∈ [0,1] which applied only to blocks of the appropriate set size.

In addition to the base models, 3 additional variants were included for each RL-only model class.

#### Choice kernel variant

First, a choice kernel was included to reflect the decaying influence of previous choices on current choice during the learning phase of the task. The choice kernel (*Ck*) acts as a weighted trace of previous actions (i.e., pressing ‘J’,’K’, or ‘L’), and is initialized to 0 for each action.

Following each trial, the choice kernel is updated for the selected action and the stored action values decay at rate *τ* according to:

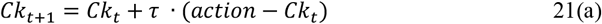

where *action* is an array of length 3 with value=1 for the chosen action and value=0 for each of the two unchosen actions on the current trial.

Each action decision is directly influenced by the choice kernel according to perseveration parameter κ:

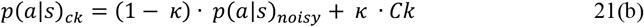

#### Forgetting variant

Next, a forgetting parameter (denoted as φ_*RL*_ to distinguish it from the working memory decay parameter φ in the RLWM models) was added to the model, which parameterized decay of RL values back to their initial (random) values according to the following:

#### Negative feedback neglect variant

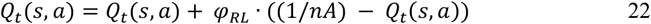

Finally, a parameter for negative feedback neglect (*η*_*RL*_) was included to allow for reduced incorporation of non-rewarded trials according to:

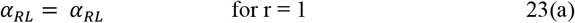

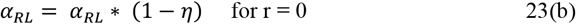

### Parameter correlation methods for additional analysis

We investigated the relationship of scores on the MASQ AA with each parameter of these 8 RL-only model variants to illustrate the potential attribution of variance by these models relating to the effects of interest from the main analysis. We used Spearman correlations corrected for FWE across all comparisons (MASQ AA x number of parameters) within each model and compared these results to the findings from the winning RLWM model (Model #5). We note that the significant effects from the winning RLWM model (Model #5) do not change in significance whether corrected for FWE across 10 comparisons (as in the main analysis) or across 9 comparisons (which would reflect correction across MASQ AA for all 9 of the model parameters). See Figure 6 for comparative results of additional analysis.

## Supplemental materials

**Figure S1.**
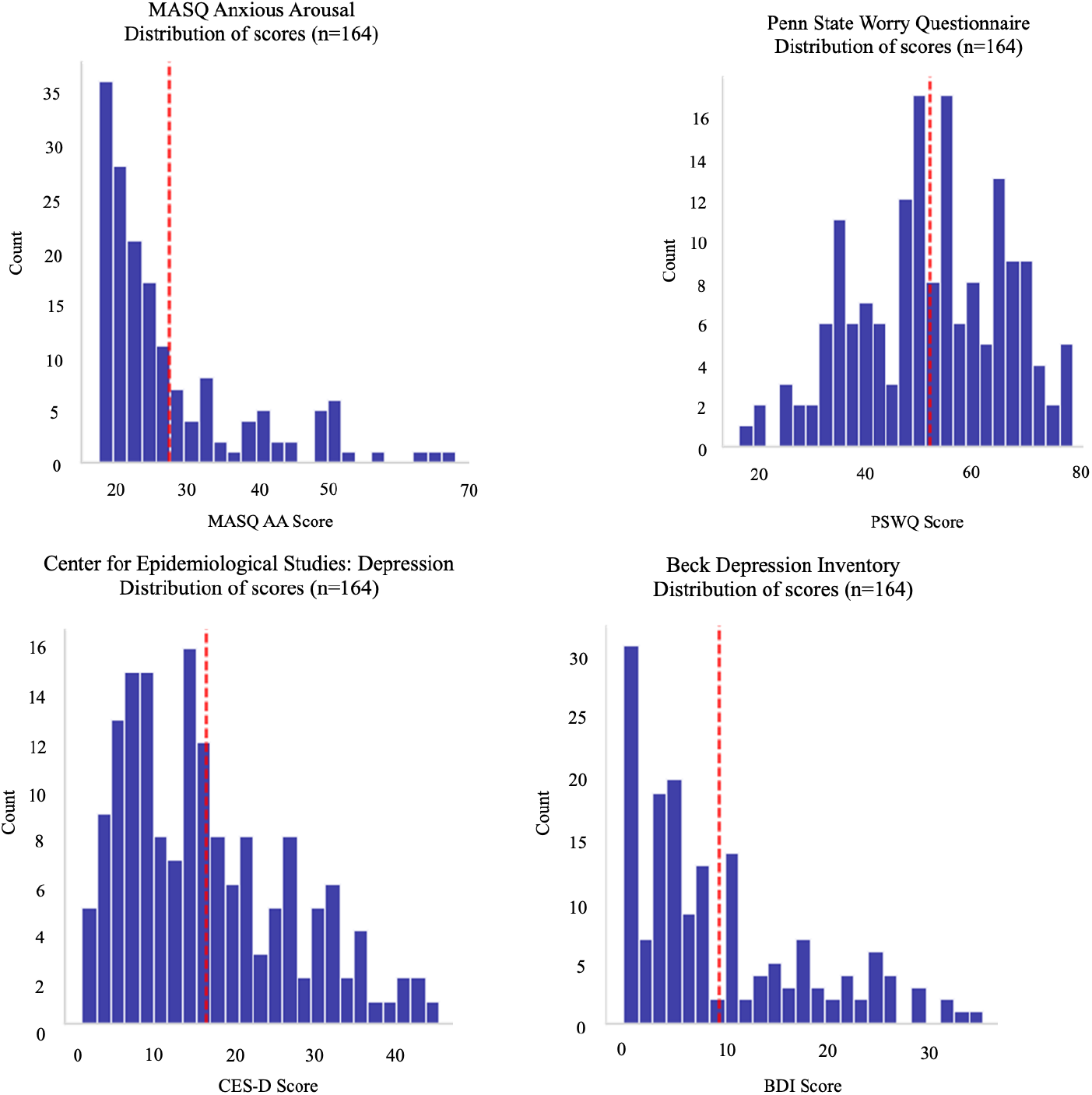
Distributions of questionnaire metrics in final participant group (n=164) Histograms showing group distribution of scores for MASQ AA, PSWQ, CES-D, and BDI-II as labeled. Red dashed vertical line denotes group mean score.

**Figure S2:**
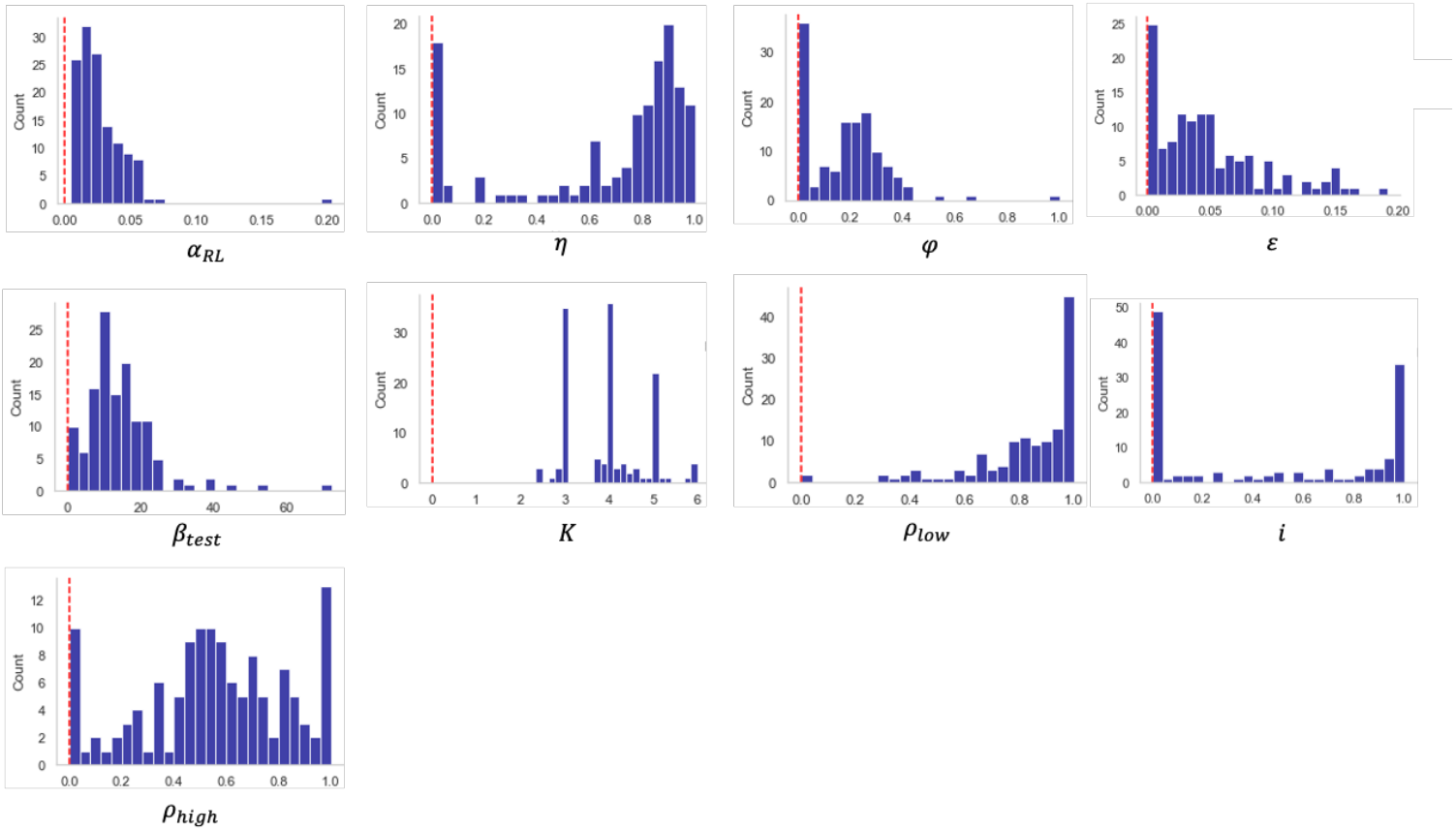
Parameter value distributions: Model #5. Histograms showing group distribution of parameter values for winning model (Model #5) as labeled. Red dashed vertical line at 0 for cross-plot comparison.

**Figure S3:**
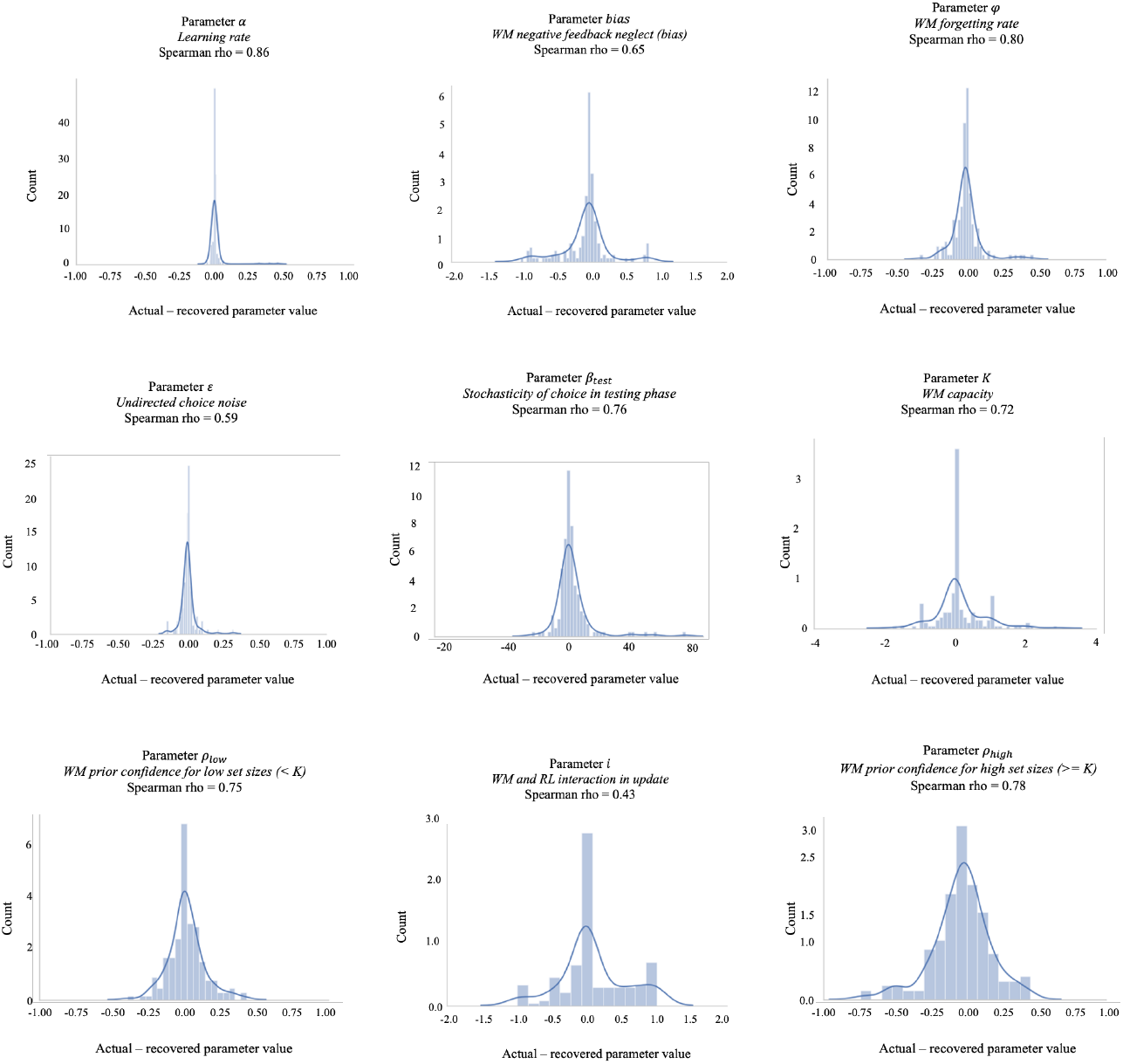
Parameter recovery analyses: Model #5. For each parameter in the winning model (Model #5), the distribution of differences between actual and recovered parameter values is shown. The correlation between actual and recovered parameter values (as measured with Spearman rho) is denoted in text above each plot. See Methods for details.

**Figure S4:**
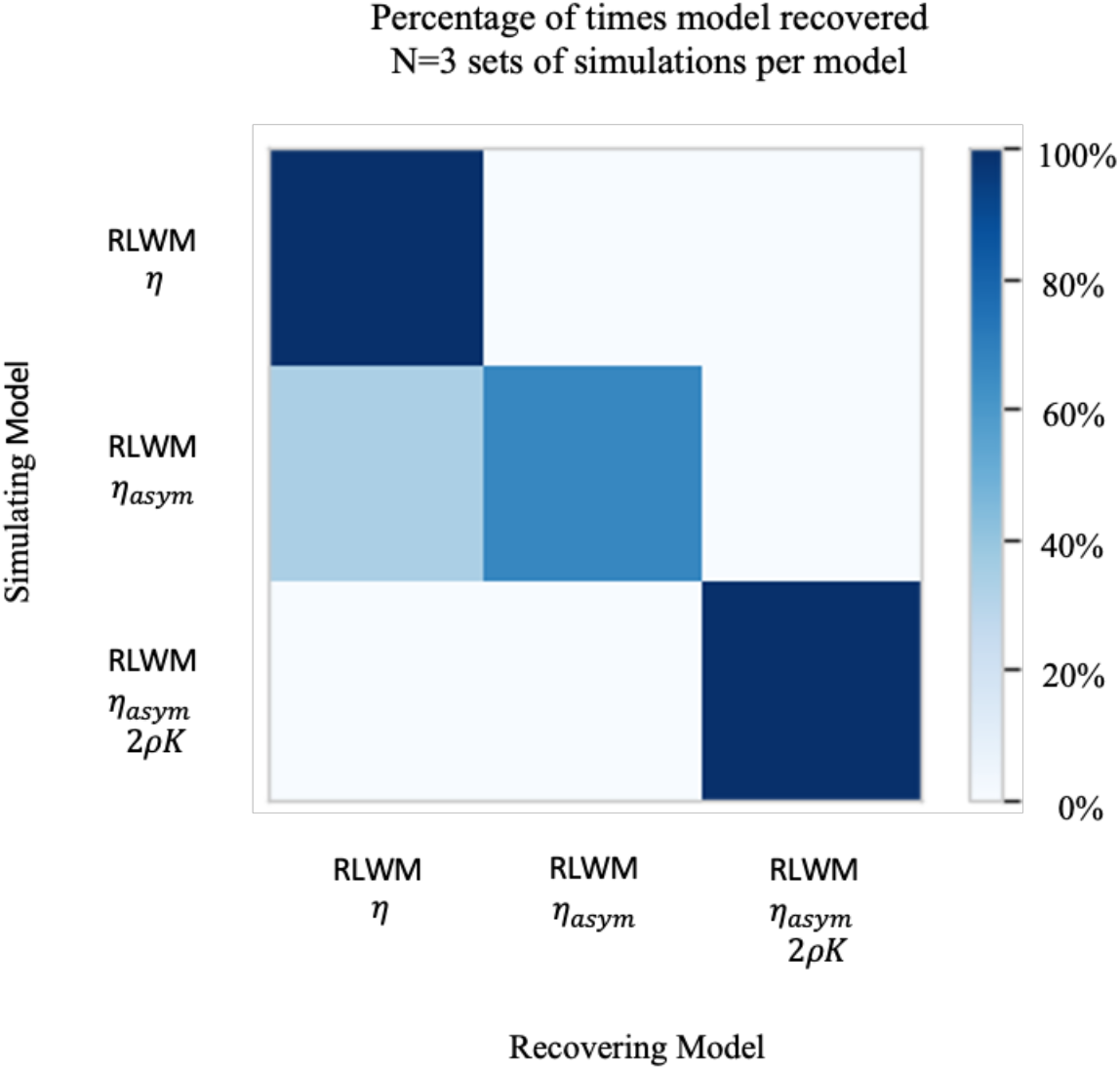
Model recovery analysis. Comparison of frequency with which the generative model provided the best fit to the data across the top 3performing models (based on AIC comparison). See Methods for details.

**Figure S5:**
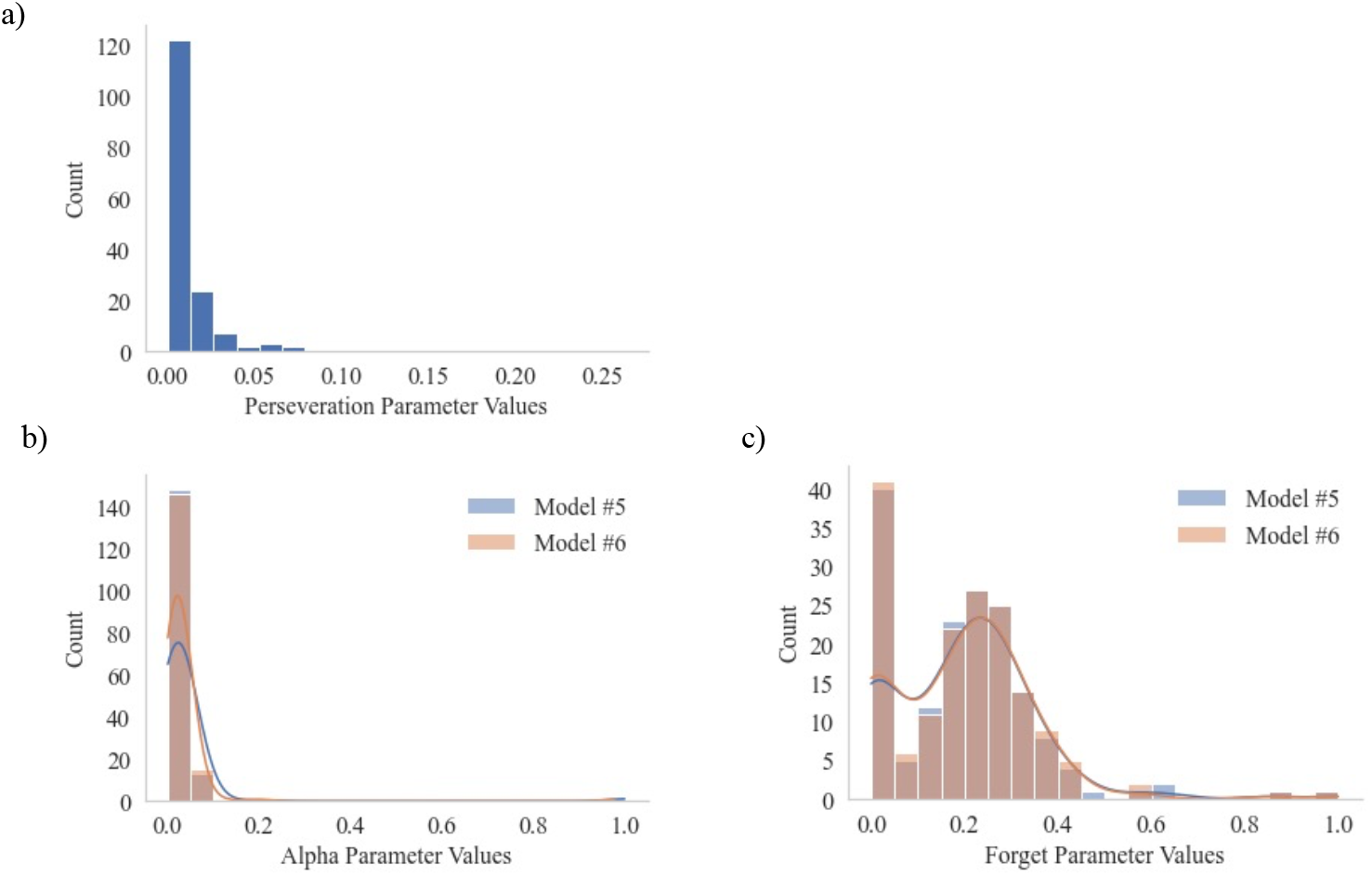
Robustness check of winning model (Model #5) against choice kernel model variant (Model #6) a) Perseveration parameter values from Model #6 reveal minimal perseveration across participants. b) Learning rate parameter values for Model #5 are not significantly different from values for Model #6 (Man-Whitney-U test: U-statistic=13,796.0, p=0.686). c) WM decay parameter values for Model #5 were not significantly different from values for Model #6 (Man-Whitney-U test: U-statistic=13,669.0, p=0.797).

**Figure S6:**
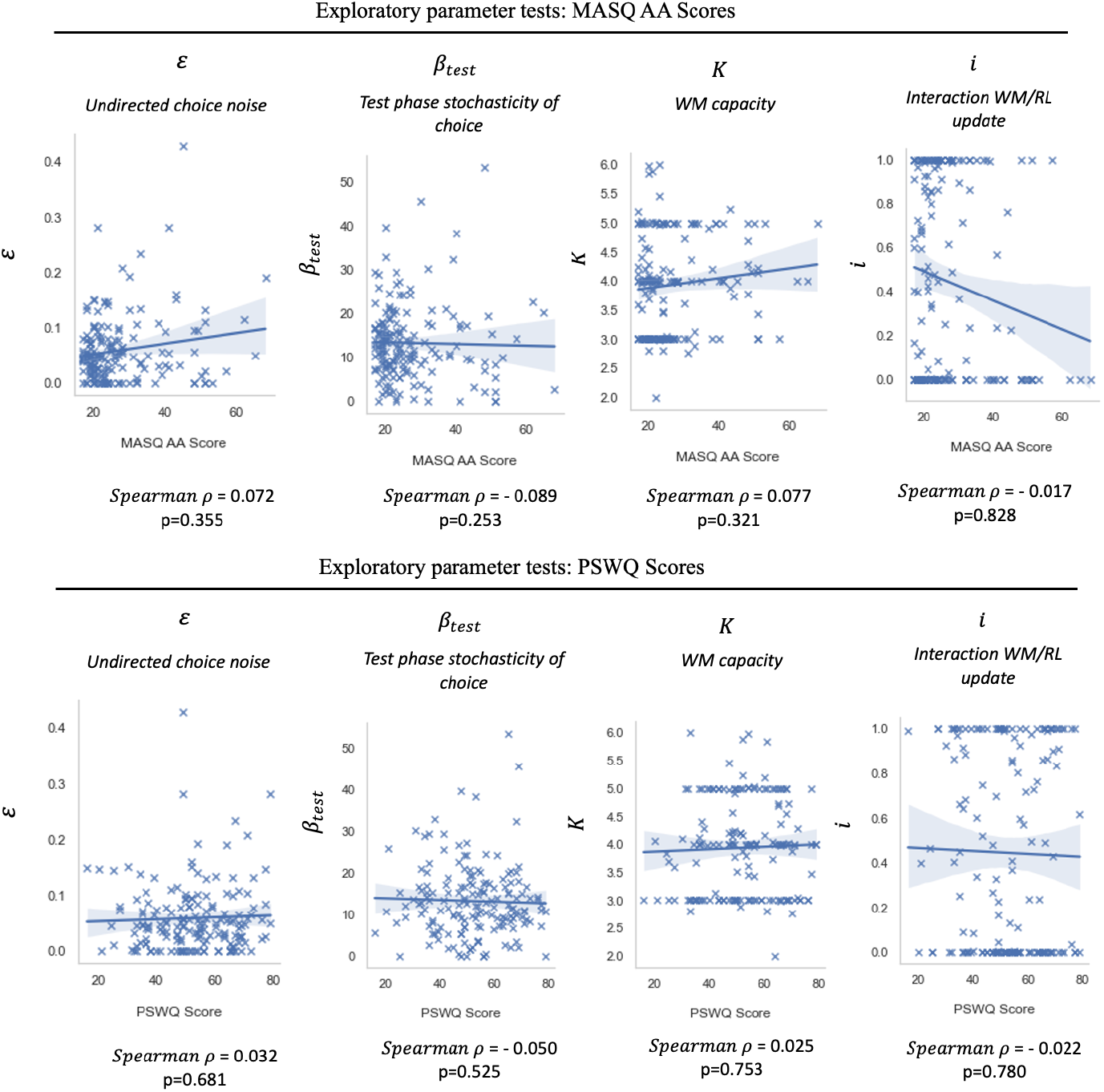
Exploratory parameter testing Model #5 for MASQ AA and PSWQ. Exploratory testing of Model #5 parameters with MASQ AA and PSWQ scores. p-values uncorrected for multiple comparisons/family-wise error.

**Figure S7:**
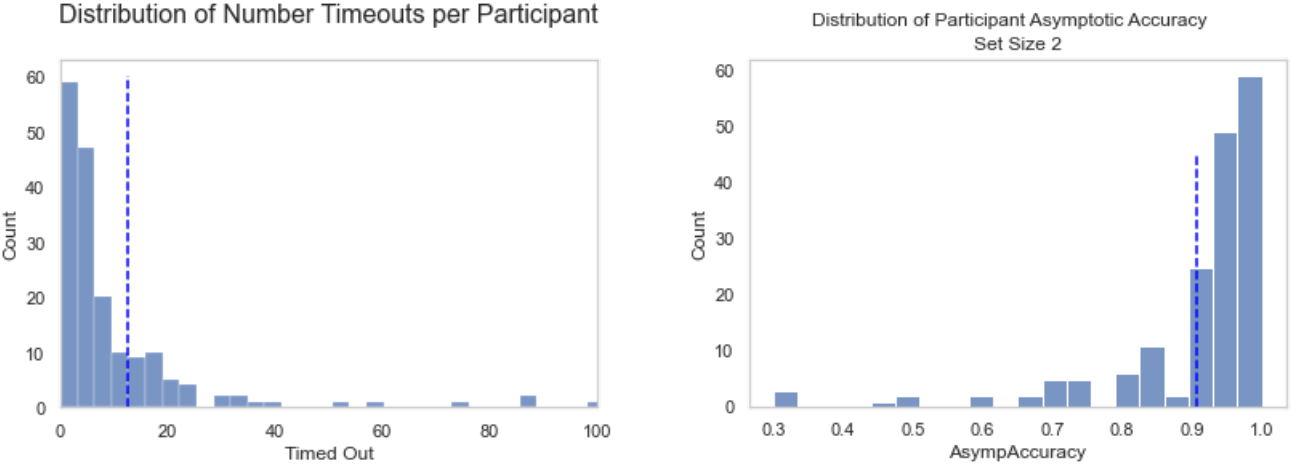
Exclusion statistics. Distribution of number of timeouts per participant prior to timeout-based exclusions (left) and distribution of participant asymptotic accuracy in learning over last 5 stimulus presentations at set size 2 (right). Dashed blue lines denote mean. Exclusion criterion was 2SD from mean (mean + 2sd for timeouts; mean – 2sd for set size 2 performance).

## Supplemental Analyses

### Test of winning RLWM model (Model #5) robustness to bias when choice kernel is included

Recent work (43) has established that in some (but not all) RL data sets, the absence of a choice perseveration kernel in the associated RL model can bias estimation of negative feedback neglect. To ensure that our winning model, which did not include a choice perseveration kernel, did not result in biased parameter estimation relevant to results, we tested the fit of the winning model (Model #5) if it was modified to include a choice kernel, allowing for a parametrically decaying influence of past choices on the current choice (Model #6; see Methods). Model #6 did not fit the data as well as Model #5 based on AIC. In addition to worse model fit based on AIC, values for the parameter quantifying perseveration were very close to 0 (see Supplements Figure 5a), and neither of the key parameters of interest from Model #5 (namely those for which there were significant findings: learning rate *α* and WM decay φ_*WM*_) differed significantly in value between Model #5 and Model #6 (see Supplements Figures 5b and 5c).

### Test of robustness of results in second-best-fitting RLWM model (Model #4)

As a robustness check across models, we tested the significant findings between MASQ AA subscale scores and model parameters in the second-closest winning model (Model #4), a nested version of the winning model with fewer parameters (notably, Model #4 did not include separate working memory confidence parameters for low versus high set sizes). Confirmatory test p-values were corrected using the Bonferroni statistical correction for 3 comparisons. As in Model #5, higher MASQ AA subscale scores were significantly related to reduced learning rate parameter (rho(162) = −0.270, FWE-corrected p = 0.004) and higher rate of working memory decay (rho(162) = 0.245, FWE-corrected p = 0.001) in Model #4. We note that if we had performed initial hypothesis testing in Model #4 instead of Model #5, we would have performed 8 total comparisons (4 model parameters from initial hypothesis testing across 2 trait measures). The relationship of MASQ AA scores with both learning rate and with rate of working memory decay in Model #4 also survive this more conservative metric of multiple comparisons (learning rate: 8-comparison FWE-corrected p = 0.003; working memory decay: 8-comparison FWE-corrected p = 0.012).

### Behavior: Reaction Time

Mean reaction time (RT) across correct learning trials was 599.5ms, and reaction times increased as a function of set size during learning (Kruskal-Wallis statistic(5) = 11308.01, p=0.0). There was a weakly trending effect of PSWQ score on longer overall mean learning reaction time (Spearman rho(162) = 0.139, p = 0.076), but no effect of MASQ AA subscale score on overall mean reaction time (Spearman rho(162) = −0.086, p = 0.276) across set sizes. A nonparametric test of correlation at each set size revealed that higher MASQ AA subscale scores were associated with a significantly shorter reaction time for set size = 5 only (Spearman rho(162) = −0.182, p = 0.02), but found no other significant relationships between MASQ AA or PSWQ scores and learning phase RTs in other set sizes (see Supplements Figure S.4.5 for plots and detailed statistics). Scores on the MASQ AA subscale were significantly related to RT set size slope in learning (Spearman rho(162) = 0.178, p = 0.023), suggesting an increased sensitivity of RT to set size increases for higher MASQ AA subscale scores. PSWQ scores were not related to RT set size slope in learning (Spearman rho(162) = −0.017, p = 0.827).

Mean RT across correct testing phase trials was 557.8ms, and was significantly lower than learning phase RT for correct trials (Mann-Whitney U statistic(1) = 617273595.0, p=1.92e-72). There was no significant effect of set size on testing phase RT (Kruskal-Wallis statistic(5) = 1.36, p=0.851). PSWQ scores were not significantly correlated with longer overall testing phase RT (Spearman rho(162) = 0.118, p = 0.131), nor were MASQ AA subscale scores (Spearman rho(162) = −0.086, p = 0.273). A nonparametric test of correlation at each set size revealed that higher PSWQ scores were associated with significantly longer reaction times for set size = 4 (Spearman rho = 0.224, p = 0.01) and set size = 5 (Spearman rho(162) = 0.174, p = 0.047) only (see Supplements Figure S.4.6 for plots and detailed statistics). There were no other significant relationships between MASQ AA or PSWQ scores and testing phase RTs by set size. RT set size slope in testing was not significantly related to either MASQ AA subscale scores (Spearman rho(162) = −0.038, p = 0.628) or PSWQ scores (Spearman rho(162) = −0.042, p = 0.595).

### Behavior: Additional statistics for PSWQ with learning and testing performance

A repeated measures ANCOVA with within-subject factor of set size, covariate of z-scored PSWQ scores, and dependent variable of mean learning performance revealed no significant main effect of PSWQ (between-subjects PSWQ F(1,162) = 1.200, p = 0.275) and no significant interaction of PSWQ and set size (within-subject set size x PSWQ effect with Greenhouse-Geisser correction F(3.415, 553.151) = 0.839, p = 0.485). There was a main within-subject effect of set size (within-subject set size effect with Greenhouse-Geisser correction F(3.415, 553.151) = 64.945, p < 0.001).

A repeated measures ANCOVA with within-subject factor of set size, covariate of z-scored PSWQ scores, and dependent variable of mean testing performance revealed no significant main effect of PSWQ (between-subjects PSWQ F(1,162) = = 0.672, p = 0.414) or interaction between set-size and PSWQ (within-subject set size x PSWQ effect with Greenhouse-Geisser correction F(3.729, 604.140) = 1.321, p = 0.262).

We additionally tested the relationship between PSWQ scores and drop between asymptotic accuracy over last 3 trials during learning and overall performance during testing. A repeated-measures ANCOVA with within-subject factor of set size, covariate of z-scored PSWQ scores, and dependent variable of drop in performance between asymptotic final 3 trial learning performance and mean overall testing performance revealed no main effect of PSWQ scores (between-subjects PSWQ F(1,162) = 0.429, p = 0.514) or interaction between set size and PSWQ (within-subject set size x PSWQ effect F(4,648) = 1.218, p = 0.302). There was a main effect of set size (within-subjects set size effect F(4,648) = 9.005, p < 0.001).

### Exploratory analyses in winning RLWM model (Model #5)

Following specific hypothesis testing, we performed an exploratory analysis of the (FWE-uncorrected) relationships between all model parameters and scores on the two measures of anxiety used for hypothesis testing above (MASQ AA and PSWQ) as well as two measures of trait depression (the CES-D scale and the Beck Depression Inventory (BDI) II). There were no significant correlations between MASQ AA, PSWQ, CESD, or BDI scores and any model parameters other than those reported for MASQ AA in the main analysis.

